# A statistical method for image-mediated association studies discovers genes and pathways associated with four brain disorders

**DOI:** 10.1101/2023.06.16.545326

**Authors:** Jingni He, Lilit Antonyan, Harold Zhu, Qing Li, David Enoma, William Zhang, Andy Liu, Bo Cao, M. Ethan MacDonald, Paul D. Arnold, Quan Long

## Abstract

Brain imaging and genomics are critical tools enabling characterization of the genetic basis of brain disorders. However, imaging large cohorts is expensive, and may be unavailable for legacy datasets used for genome-wide association studies (GWAS). Using an integrated feature selection/aggregation model, we developed Image-Mediated Association Study (IMAS), which utilizes borrowed imaging/genomics data to conduct association mapping in legacy GWAS cohorts. By leveraging the UK Biobank image-derived phenotypes (IDPs), IMAS discovered genetic bases underlying four neuropsychiatric disorders and verified them by analyzing annotations, pathways, and expression Quantitative Trait Loci (eQTLs). A cerebellar-mediated mechanism was identified to be common to the four disorders. Simulations show that, if the goal is identifying genetic risk, IMAS is more powerful than a hypothetical protocol in which the imaging results were available in the GWAS dataset. This implies the feasibility of reanalyzing legacy GWAS datasets without conducting additional imaging, yielding cost-savings for integrated analysis of genetics and imaging.

## INTRODUCTION

Brain imaging plays a critical role in neuroscience and psychiatry. Neuroimaging endophenotypes are intermediate traits closer to the action of the genes than observable phenotypes. Therefore, neuroimaging could be leveraged to identify risk genes and associated physiological processes in the brain. Projects focusing on the genetic basis of brain images, i.e., imaging genetics, have been launched and led to discoveries via associating single nucleotide variants (SNVs) to image features relevant to neurological or psychiatric disorders^1–3^. Neuroimaging has been extensively studied as a potential biomarker in diagnostics and risk assessment of neuropsychiatric disorders^4–6^. Thus, there are potential mediating roles that imaging can serve in accurately discovering the genetic basis of brain disorders.

However, neuroimaging methods, such as magnetic resonance imaging (MRI), are expensive, preventing broad utilization of neuroimaging to further characterize the genetic basis of brain disorders. Moreover, neuroimaging is unavailable for most legacy cohorts where genotype and phenotype are in place (e.g., most GWAS datasets).

Fortunately, however, a number of datasets providing brain images and genotype are available^7–9^. If researchers can redirect such existing data as a “reference panel” into their dedicated studies for preliminary discovery of putative associations, it will bring enormous cost-savings for scientific discovery as well as timely judgement in a clinic setting. The UK Biobank is a flagship database that contains genomics data for half a million individuals, including nearly 50,000 assessed for brain images^7^. Those brain images include a wealth of contrast types, including: T1 and T2 weighted, Fluid Attenuated Inversion Recovery, resting and task-based functional MRI, and diffusion MRI. Moreover, it also offers hundreds of neuroimaging endophenotypes, called image-derived phenotypes (IDPs), which are quantitative indicators representing brain structure and function. Examples of IDPs include the volume of grey matter in specific brain regions, and measures of functional and structural connectivity.

To seamlessly integrate general resources of the brain imaging and GWAS datasets dedicated to particular disorders, we developed a statistical framework, Image-Mediated Association Study (IMAS), to discover the genetic basis of brain disorders through the mediation of imaging, without assuming the images are assessed in the GWAS cohort. This work is partly inspired by the popular Transcriptome-Wide Association Study (TWAS)^10–16^, however, built upon our unique “data-bridge” angle of disentangling TWAS into feature selection and feature aggregation^17–19^. Although IMAS utilizes a “borrowed” reference panel of images instead of requiring imaging in the cohort with genotype and phenotype, it could be even more powerful than applying a naïve association on the data with both images and phenotype. This is because, assuming the genetic component is the causal factor underlying brain morphology, aggregated SNVs selected by “borrowed” images contains only the genetic component, whereas the real images suffer from noise, motion, or other factors. This insight was demonstrated in another similar setting for transcriptome-wide association study (TWAS) by us^19^. Intuitively, this effect for IMAS should be more pronounced due to the higher heritability of images in contrast to gene expressions used in TWAS. Herein, we conducted simulations using various genetic architecture parameterizations to verify this advantage for IMAS, which allows cost-savings in integrated analysis of genetics, imaging, and brain disorders.

To verify the expected advantage of IMAS over conventional protocols, we conducted analysis using IDPs from the UK Biobank and genotype/phenotype data from four neuropsychiatric disorders, i.e., schizophrenia (SCZ)^20^, major depression disorder (MDD)^21^, bipolar disorder (BPD)^22^, and autism spectrum disorder (ASD)^23^. We observed that: (1) IMAS identified IDPs that are relevant to corresponding neuropsychiatric disorders; (2) the underlying SNVs supporting the associations between IDPs and disorders are enriched in sensible pathways; and (3) the top SNVs are eQTLs (expression quantitative trait loci) that regulate critical genes in the corresponding conditions. Notably, a cerebellar-mediated mechanism was identified to be shared in the four disorders. This finding is consistent with recent emerging evidence that cerebellar abnormalities may confer broad risk to psychopathology across disorders^24^.

## RESULTS

### The IMAS model and implementation

IMAS is partly inspired by the method of transcriptome-wide association studies (TWAS) that have been extensively developed^10–16^ and applied^16, 25–34^ recently. The mainstream format of TWAS is a two-step protocol: First, one trains an expression prediction model using (cis-) genotype: *E* ∼ Σ β_i_X_i(Ref)_ + ∈ in a reference dataset, e.g., GTEx^35^, in which the expression and genotype are available (**Figure 1A**). The predicted expression is called Genetically Regulated eXpression (GReX). Second, in the main dataset for GWAS (that doesn’t contain expression), one can predict expression using the genotype: *Ê* = Σ β_i_X_i(GWAS)_, and then use the predicted expression, *Ê*, to conduct the association test (**Figure 1B**). The outcome will be the gene associated to the trait; and this may be applied to all genes with heritability, e.g., >1%.

**Figure 1.**
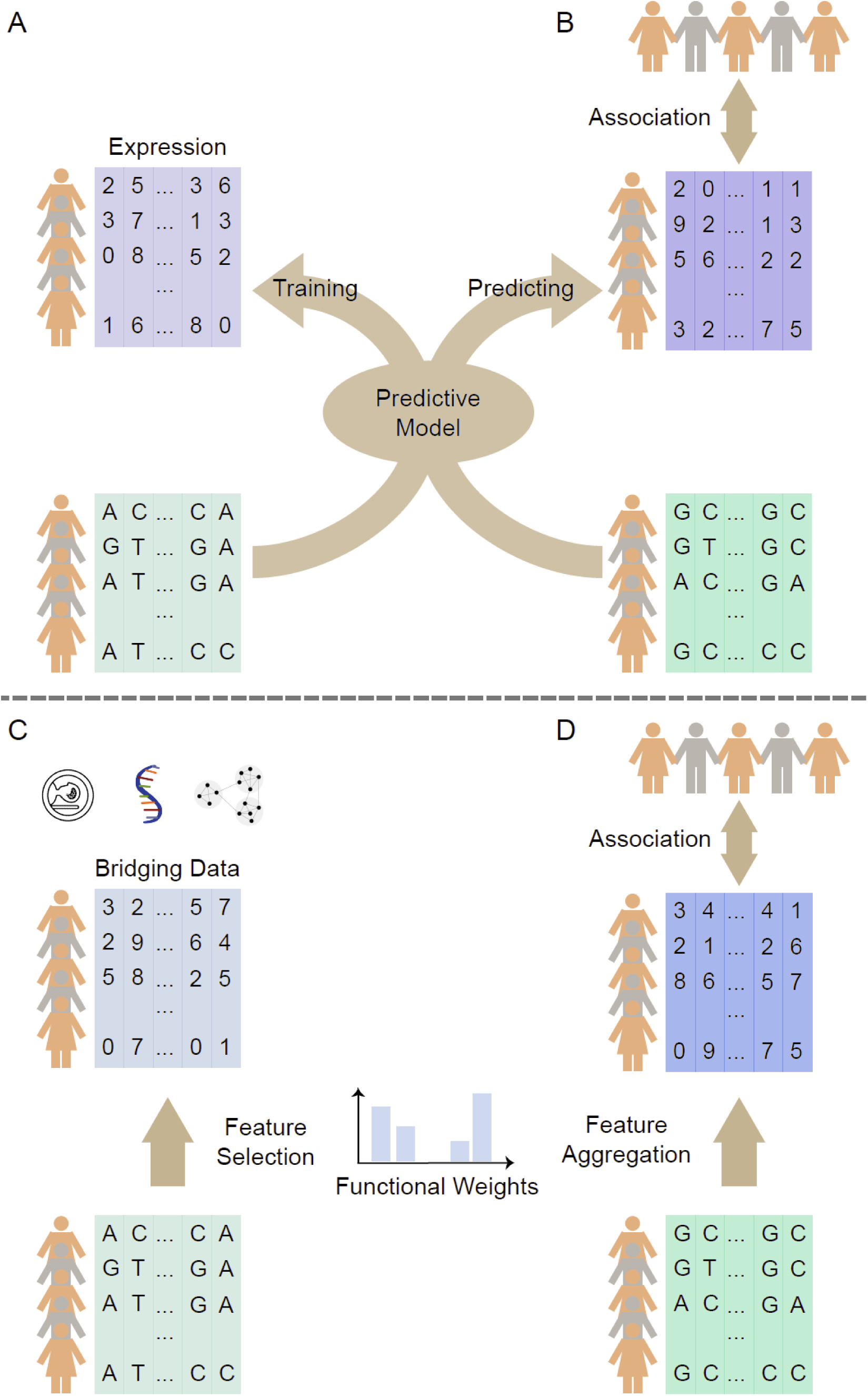
The data-bridge framework in IMAS. Standard TWAS (A and B) and the *data-bridge* Framework adapted in IMAS (C and D). A. Standard TWAS uses a reference dataset to train a predictive model of gene expression using genotype data. B. This predictive model is then used to predict expressions in the GWAS dataset (where expression is unavailable). The predicted expression is associated with the phenotype of interest. C. The *data-bridge* framework uses a reference dataset (e.g., IDPs used in IMAS) for the selection of genetic features and their weights. D. Weights derived from the feature selection step are then used in feature aggregation for the GWAS dataset. These aggregated features are then associated with the phenotype of interest to identify variants of interest. Both TWAS and IMAS are special cases of the data-bridge framework.

Our recent mathematical analysis^17–19^ indicated that “predicting expression” does not represent the essence of TWAS and may be misleading. As such, we proposed a “data-bridge” framework to interpret the “prediction” step as the selection of genetic variants (and their functional weights) directed by expression (**Figure 1C**) and interpret the use of predicted expression in the second step as feature aggregation (**Figure 1D**) by a linear model. These are two distinct steps in Machine Learning. With this interpretation, the original TWAS dubiously binds two distinct steps into the same statistical form (GReX). Instead, if we split them and consider gene expressions as a “data-bridge”, methodological research focusing on optimal combinations of feature selection and aggregation approaches will be more powerful. Consistent with this insight, alternative methods splitting these two steps developed by us^17, 18^ and independently by others^36^ are superior to standard TWAS.

Enabled by this framework, IMAS considers reference images as a data-bridge to link genotype and phenotype. From a dataset with imaging assessment (such as IDPs in the UK Biobank) and genotype, IMAS learns SNVs that are functionally relevant to images. Then, for a GWAS dataset with genotype and focal disease phenotype, IMAS aggregates selected genetic variants to determine the association of each aggregated SNV-set with the phenotype, reporting the association between the genetic component and the risk of disease. Particularly, IMAS provides two optional models for feature selection: (1.a) elastic-net, a regularized multiple-regression^10^ the default method in TWAS, and (1.b) a test for each SNV individually (i.e., testing for the marginal effect) using linear mixed model^18^. For the feature aggregation step, there are also two optional models: (2.a) the linear combination of selected SNVs, and (2.b) the kernel machine as implemented by SKAT^37^. The above models form 2 x 2 = 4 combinations of protocols offered by IMAS (**Online Methods**). Based on our previous comparison^17, 18^ and that of others^36^, kernel methods are more powerful than linear combinations in aggregating signals. Additionally, our previous research shows marginal effect-based selection outperforms elastic-net in most cases^18^. This is also confirmed by the IMAS-tailored simulations, as will be presented. Therefore, the recommended IMAS-protocol is (1.b) + (2.b); and the real data analysis in this paper follows this recommendation. The software is freely available at our GitHub (**Web resources**).

### IMAS discovered IDPs associated with neuropsychiatric disorders

We applied IMAS to the GWAS datasets (i.e., genotyping data and disease labels without neuroimaging) of four brain disorders, i.e., schizophrenia (SCZ), major depression disorder (MDD), autism spectrum disorder (ASD) and bipolar disorder (BPD). By utilizing IDPs “borrowed” from the UK Biobank, IMAS was able to identify three, eight, fourteen and one significant IDP(s) that are associated with SCZ, MDD, ASD, and BPD respectively (based on a Bonferroni-corrected P-value < 0.05) (**Table 1; Figure 2A,D,G,J**). We conducted literature searches on the significant IDPs (used the 3 and 1 significant ones for SCZ and BPD, respectively, and selected the top four for MDD and ASD) in relation to their corresponding disorders. We were able to annotate the associations in the context of extensive existing work on neuroimaging studies (**Supplementary Notes I**).

**Figure 2.**
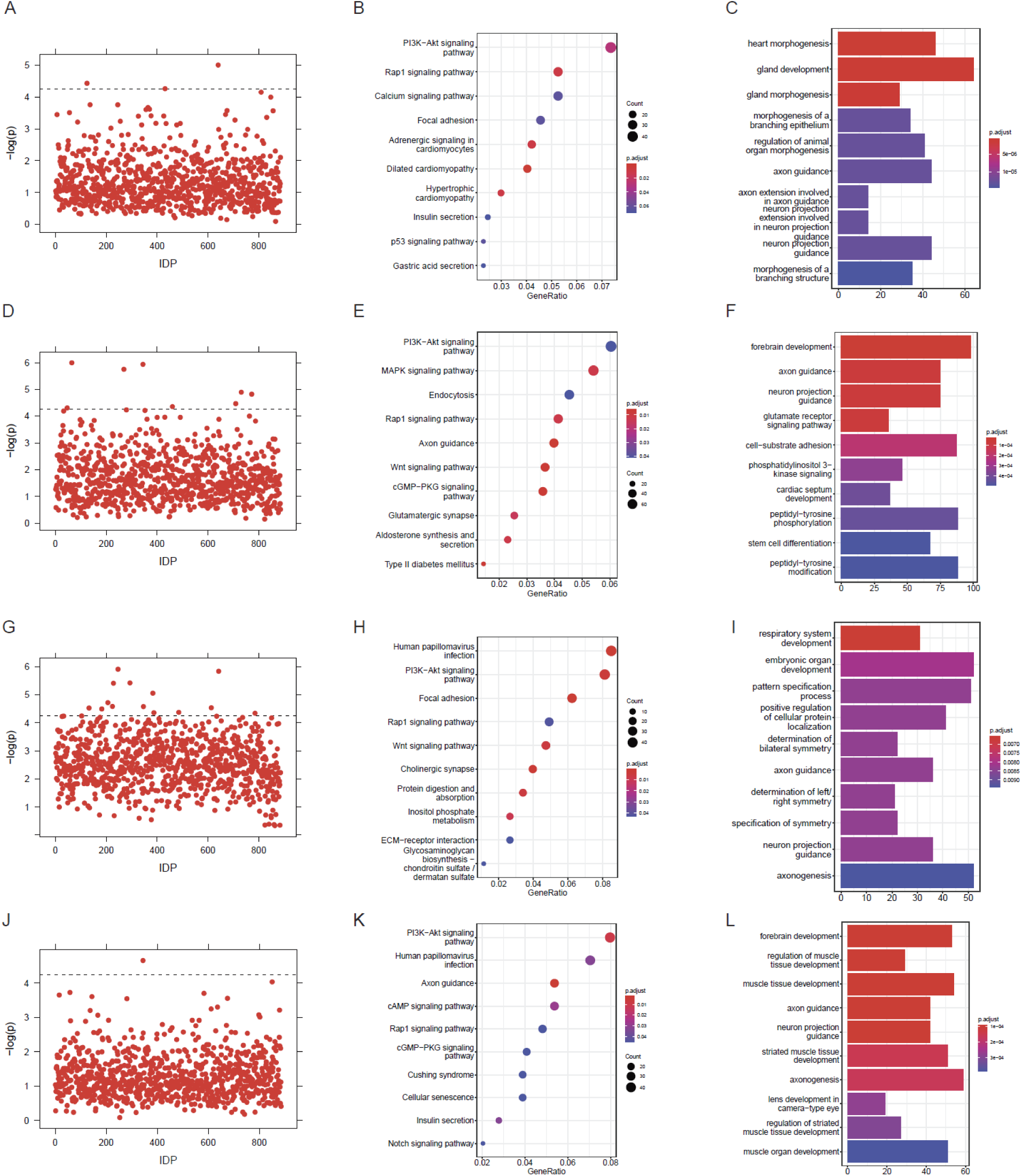
Applying IMAS to four neuropsychiatric disorders. A,D,G,J. Dashed line indicates the cutoffs of Bonferroni-correction (= 0.05/total number of IDPs). A. Schizophrenia. D. Major Depression Disorder. G. Autism Spectrum Disorder. J. Bipolar Disorder. B,E,H,K. The KEGG pathway analysis for IMAS identified SNPs. We selected the most significant IDP for each disorder to present. All the panels are dot plot showing the top 10 KEGG pathways ranked by the GeneRatio values. The size of the balls indicates the number of the genes enriched and the color indicates the level of the enrichment (P-adjusted values). The GeneRatio is calculated as count/setSize. ‘count’ is the number of genes that belong to a given gene-set, while ‘setSize’ is the total number of genes in the gene-set. C,F,I,L. The GO enrichment analysis for IMAS identified SNPs for four neuropsychiatric disorders. All the panels are bar plot showing the top 10 enriched biological processes ranked by p-values. The correlation is more significant as the red/blue ratio increases. The number on the x-axis indicates the number of genes that belongs to a given gene-set. B.C. Schizophrenia, IDP-25190-2.0. E.F. Major Depression Disorder, IDP-25152-2.0. H.I. Autism Spectrum Disorder, IDP-25869-2.0. K.L. Bipolar Disorder, IDP-25904-2.0.

**Table 1.**
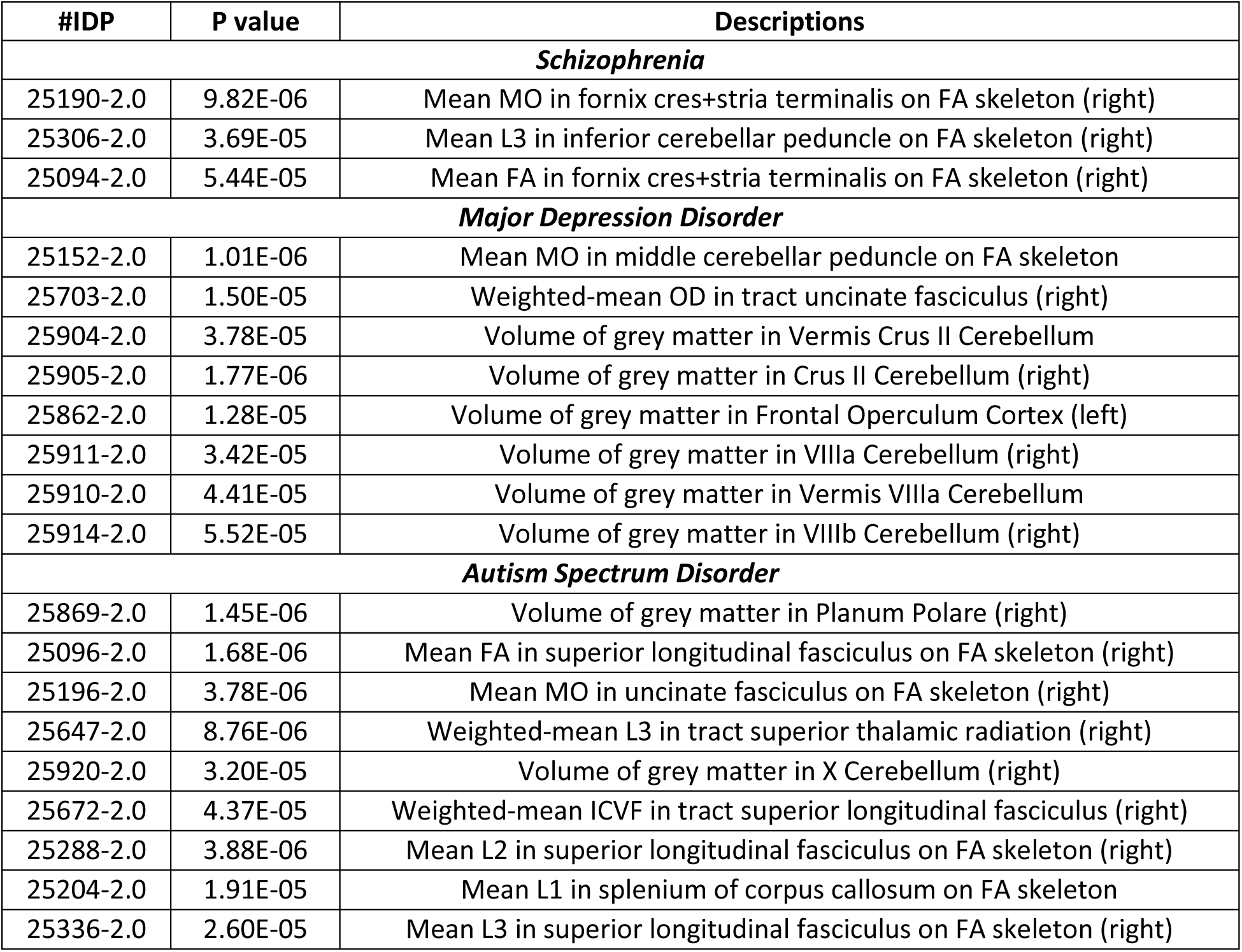
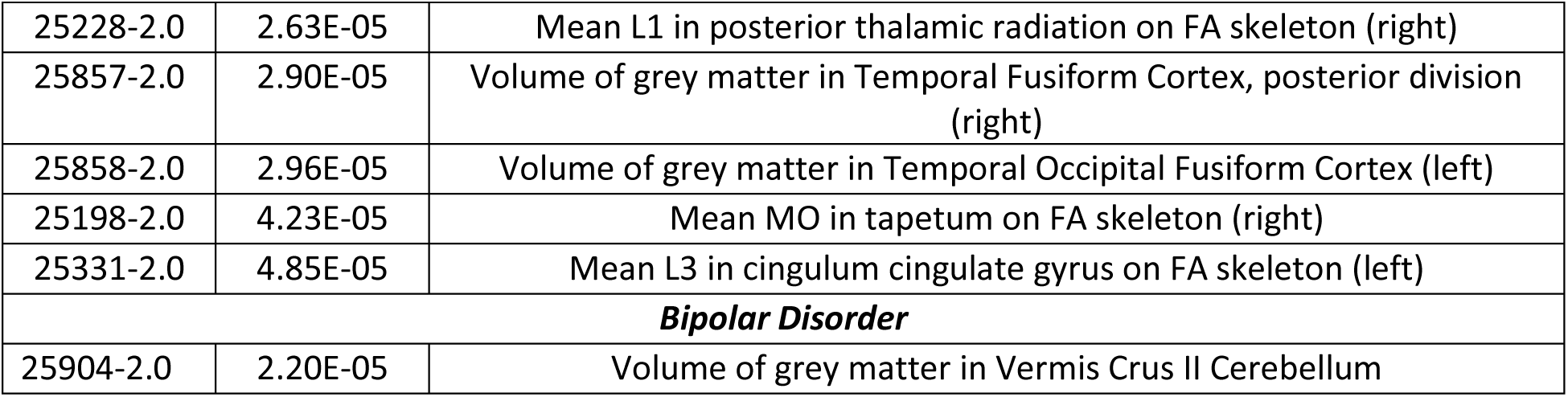
Significant IDPs and their corresponding *P* value and descriptions for schizophrenia (SCZ), major depression disorder (MDD), autism spectrum disorder (ASD), and bipolar disorder (BPD).

### IMAS identified SNVs enriched in sensible pathways

The above discovery of IDPs is supported by the underlying putative genetic variants that are supposed to be associated with corresponding disorders. We then further conducted GO enrichment and KEGG pathway analysis towards the genes where the SNV sets are located (by mapping each SNV to its “functionally closest” gene (**Online Methods**). We showed that SNV-sets identified by IMAS are indeed enriched in sensible pathways in terms of both KEGG and GO analysis. The KEGG and GO pathways for the gene sets associated with the most significant IDP in each disorder are displayed in **Figure 2B,E,H,K**, and **Figure 2C,F,I,L**, respectively. The KEGG pathways for the rest of the IDPs are listed in **Supplementary Table 1**, and the GO enrichment of the rest IDPs are listed in **Supplementary Table 2, 3, and 4**, listing Biological Processes, Molecular Function, and Cellular Component, respectively. Interestingly, the PI3K/AKT pathway appears in all disorders. We further examined the experimental evidence to show the relevance of these pathways and GO terms in relation to the corresponding disorders and indeed discovered massive evidence for all the top five sets (**Supplementary Notes II**).

### The top SNVs overlapped with eQTLs that regulate genes functionally important for neuropsychiatric disorders

To link the IDPs to gene regulatory mechanisms, we conducted eQTL analysis based on GTEx-reported eQTLs^38, 39^ (**Online Methods**). We required a strict criterion that identical locations of the IDP-selected SNVs and eQTL should be matched. As a result, many associated SNVs are not mappable to the eQTLs, with the proportion of successfully mapped IMAS SNVs (to a GTEx SNV that is an eQTL) ranging from 11.9% to 96.6% with a mean of 22.2% (**Supplementary Table 5**). Among these mapped eQTL SNVs, we chose to analyze the top five for each IDP, and indeed verified the critical roles of these genes in the pathology of the related disorders. The detailed descriptions of literature supports are in **Supplementary Notes III** and the regulatory networks are depicted in **Figure 3**.

**Figure 3.**
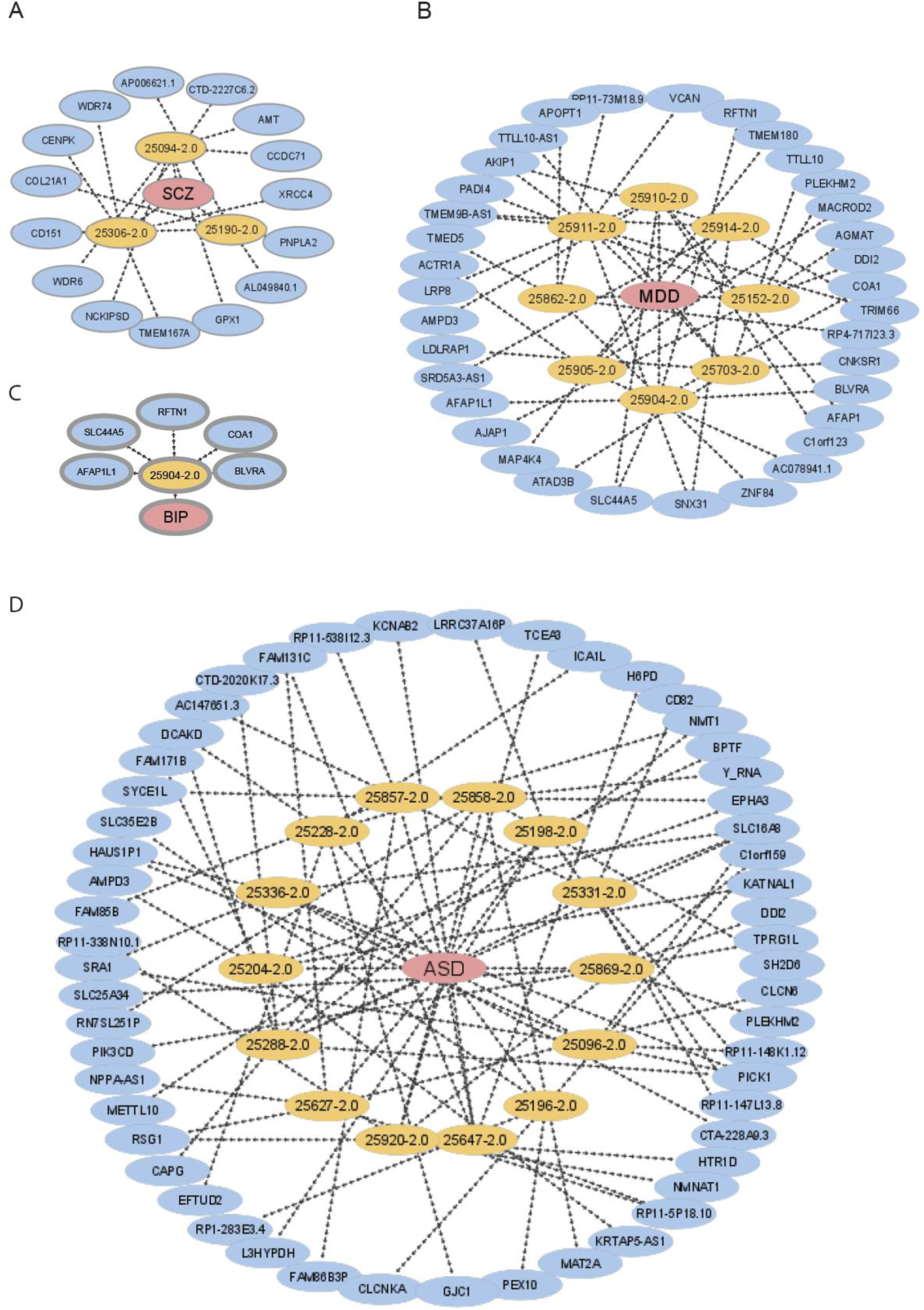
Top IMAS-discovered SNVs overlap with eQTLs regulating functionally relevant genes for neuropsychiatric disorders. Red ovals indicate four neuropsychiatric disorders. Yellow ovals are significant IDPs for each disorder. Blue ovals are top five eQTLs regulated genes. A. Schizophrenia. B. Major Depression Disorder. C. Bipolar Disorder. D. Autism Spectrum Disorder.

### Standard GWAS cannot discover the IMAS-selected SNVs and pathways

To compare IMAS with standard GWAS, we used linear mixed model (LMM)^40, 41^ to identify genetic variants associated with the susceptibility to the four neuropsychiatric disorders (**Supplementary Figures 1-4**). Upon Bonferroni correction, except for ASD, standard GWAS methods did not discover any significant variants. With a less stringent P-value cutoff of 10^-5^, we were able to identify some variants. We conducted GO enrichment and KEGG pathway analysis using the same pipeline for IMAS outcomes (**Supplementary Table 6**). Evidently, the enriched pathways do not show clear evidence associated with the related disorders. For instance, in ASD (the only one with significant SNVs using Bonferroni correction), the enrichment is mostly related to antigen processing, which have some sort of indirect links with ASD, but not as well-known as brain functions, such as neurotransmission, synaptic dysfunction, neuron recognition and neurogenesis^42–44^. At the loose threshold of selecting SNPs of 10^-5^, there are no enriched GO terms being identified for ASD. For another example, for SCZ, most GO terms are enriched in processes related to cell-substrate adhesion. There are some literatures stated that focal adhesion dynamics are altered in SCZ^45^, and cell adhesion molecules help determine the systems-level structure of the nervous system during development and regeneration^45^, but none of the pathways that were identified by our methods appeared to be enriched. The results for KEGG pathway analysis also did not identify any pathways that were identified using our methods (**Supplementary Table 7).**

### A genetic mechanism underlying cerebellar abnormalities shared by four disorders

Numerous theories have been proposed to explain the pathophysiology of psychiatric disorders, specifically SCZ, MDD, ASD and BPD. Utilizing the IMAS approach new theories can be delineated that integrate data derived from the IMAS tool with evidence from existing literature.

Across the whole brain IDPs, one brain region emerged as being particularly noteworthy. Significant abnormalities were identified in the cerebellum IDPs in all four disorders: grey matter changes in MDD (IDP-25904-2.0, IDP-25905-2.0, IDP-25911-2.0, IDP-25910-2.0, IDP-25914-2.0), ASD (IDP-25920-2.0) and BPD (IDP-25904-2.0); white matter changes in SCZ (IDP-25306-2.0) (**Table 1**; **Figure 4A**).

**Figure 4.**
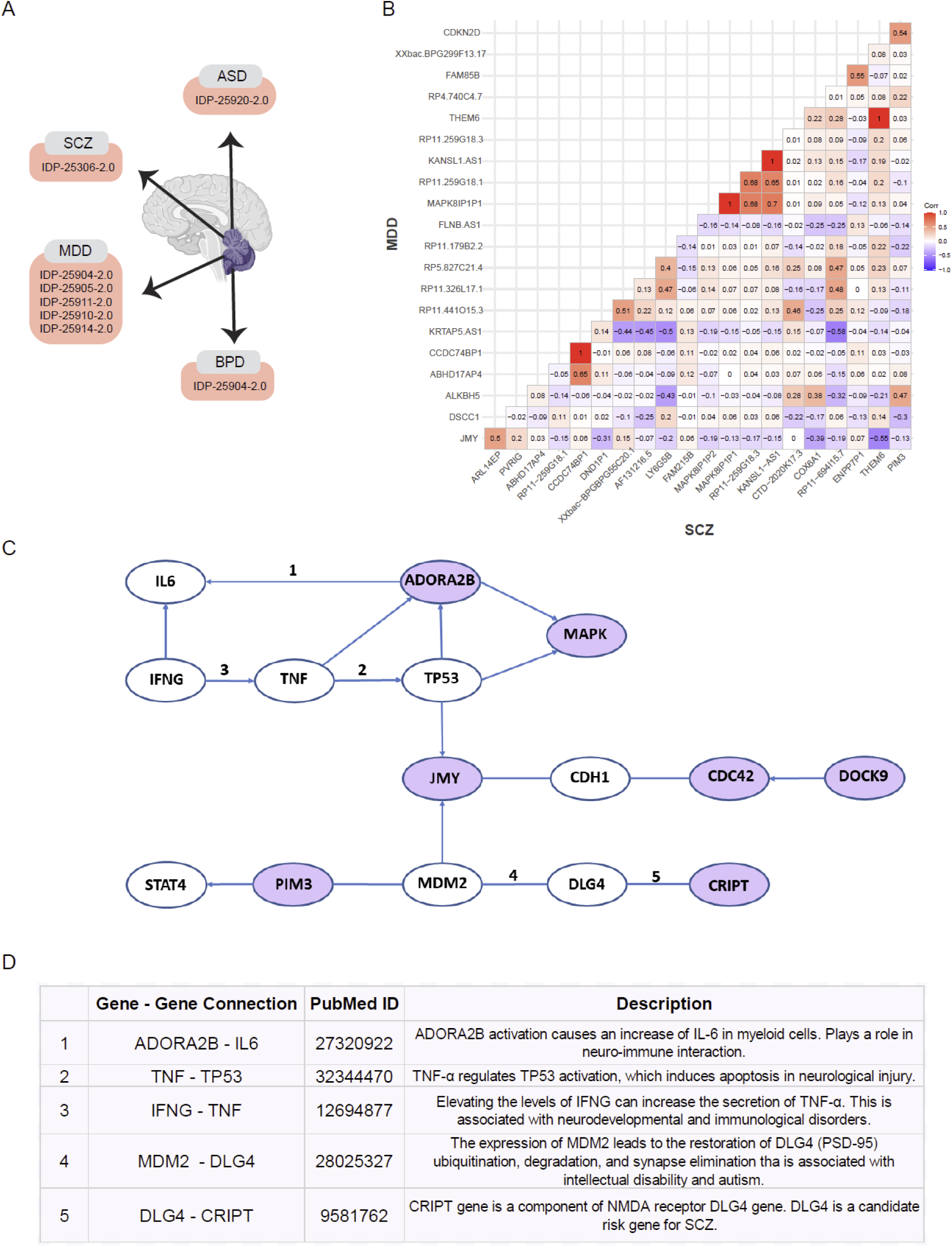
IMAS identifies underlying genetic mechanism correlated with cerebellum for four main neuropsychiatric disorders. A. An illustration of cerebellum IDPs that were identified by IMAS in all four disorders: schizophrenia (SCZ), major depression disorder (MDD), autism spectrum disorder (ASD) and bipolar disorder (BPD). B. A cross-disorder co-expression matrix between the expression level of cerebellum eQTLs that were identified from MDD and SCZ. C. A gene to gene connection network includes gene-gene interactions that are identified by literature-based analysis of publications. Nodes are genes. Links are gene-gene interactions. A link with an arrow shows regulation and the direction, a non-arrow link shows reported correlation. Purple nodes are genes that were identified from our analysis. White nodes were identified from the literature, and they connect the genes identified from our analysis. All the gene-gene interactions have been reported in literature. Numbers on the lines are “Gene-Gene Connections” mentioned in the table below. D. Five connections are selected as they play crucial roles in neurodevelopmental and psychiatric pathophysiology according to this mechanism.

To delineate the shared genetic mechanism of the four mental disorders, we followed up on the associations with cerebellum by expanding to an underlying network identified via knowledge-based analysis. For a given pair of disorders, e.g., MDD and ASD, we calculated the co-expression matrix of IMAS identified MDD-regulating genes and ASD-regulating genes in the GTEx cerebellum tissue (**Online Methods**). (**Figure 4B; Supplementary Figures 5-9**).

As shown in **Figure 4C**, various genes in our co-expression matrix have direct and/or indirect relationships with *ADORA2B* (adenosine A2b receptor). *ADORA2B* is involved in axon elongation through its interactions with *netrin-1*^46^. It plays a crucial role in brain aging^47^ and in general cognitive function^48^. *ADORA2* receptors have been identified as a risk gene for Lesch-Nyhan syndrome, a congenital disorder that severely affects the brain and behaviour of children^49, 50^. *ADORA2B* also interacts in *MAPK* pathway and associated signals^51, 52^ (**Figure 4C**). In our study, a long non-coding region of the *MAPK* gene is expressed in cerebellum IDPs of MDD, SCZ and ASD (**Figure 4B; Supplementary Figures 5-9; Supplementary Table 8**). Further, TP53 gene controls *MAPK* signalling via *Ras1/Raf1* kinase activation^52–54^. Shao et al. show that in neurological injury *TP53* activation is regulated by *TNF-α* and induces apoptosis^55^ (**Figure 4C)**. If *IFNG* gene levels are elevated, TNF increases in individuals with neurodevelopmental and immunological disorders^56^ (**Figure 4C)**. *IFNG* controls IL6 gene regulation^57^ which can be increased in myeloid cells if *ADORA2B* is activated^58^. This sub-network plays a crucial role in neuro-immune interaction which has been proposed as a potential mechanism in many neurodevelopmental and psychiatric disorders^58–60^ **(Supplementary Table 9).**

Furthermore, *TP53* interacts with *ADORA2B* and also controls the encoding of *JMY* (junction-mediating and regulatory) protein^61^ (**Figure 4C)**. *JMY* is related to cytoskeleton dynamics and upregulates myelination in actin polymerization and oligodendrocyte differentiation^62^. It also plays a major role in cortical development^63^. We show that JMY is negatively correlated with *DOCK9* (corr = −0.35 and co-expressed in cerebellum IDPs of MDD, SCZ, ASD and BPD (**Figure 4B; Supplementary Fig 5-9; Supplementary Table 8**). Moreover, *CDH1* (epithelial cadherin) and *JMY* expression leads to downregulation of adhesion molecules of the cadherin family^64–66^(**Supplementary Table 9**). Further, *DOCK9* gene enables cadherin binding activity^68, 69^ and is predicted to be involved in positive regulation of an activator of Rho-GTPase Cdc42. *DOCK9* has been associated with BPD with regards to both risk and increased illness severity^67^. *Cdc42* in Bergmann glia was found to play an important role during the late phase of radial migration of cerebellar granule neurons, important in cerebellar corticogenesis^68^. According to the co-expression matrices DOCK9 is also negatively correlated with CRIPT (corr = −0.46) and co-expressed in cerebellum IDPs of MDD, ASD and BPD (**Supplementary Fig 5-9; Supplementary Table 8**). *CRIPT* is a component of NMDA receptor DLG4 (alternative gene name is PSD-95) complex^69^. It plays a major role in synaptogenesis and synaptic plasticity^70^ and is a risk candidate gene for SCZ^71, 72^. Further, the expression of *MDM2* leads to the restoration of *PSD-95* ubiquitination, degradation, and synapse elimination in the neurons of a mouse model of Fragile X Syndrome, a leading genetic cause of intellectual disability and autism^73^. *MDM2* is also involved in the regulation of JMY protein state changes and *STAT4* gene regulation via expression of *PIM3* activation^74^ (**Supplementary Table 9**).

Overall, we propose based on our IMAS findings that, consistent with previous literature, the cerebellum plays a pivotal role in mediating cross-disorder genetic effects on multiple neuropsychiatric disorders. Based on our findings, the mechanism underlying cerebellar alterations across disorders may relate to pathways previously implicated in psychiatric genetics, such as immune inflammation pathways, and synaptogenesis.

### Simulations show that IMAS outperforms direct association mapping between actual IDPs and disorders

The above real data analysis “borrows” the UK Biobank IDPs to re-analyze GWAS data. Here, by simulations, we demonstrate that even if the IDPs are in place for the legacy GWAS datasets, the statistical power of IMAS is higher than a naïve association mapping between images (IDPs) and the phenotype, which is called “Direct association” hereafter (**Online Methods**).

We first checked the type I errors of IMAS. For all the four protocols, the 5% cutoffs (determined by simulating traits under the null distribution (**Online Methods**)) are around 0.05 (**Supplementary Table 10**). So, the type I error of IMAS is under control.

Assuming that actual images are available in the GWAS datasets, we tested power of IMAS under both pleiotropy and causality architectures in contrast to Direct-association (Di-ASSO) based on linear genetic architectures (**Online Methods**). We estimated heritability of each IDP using GCTA^75^ and the UK Biobank dataset (**Supplementary Figure 10**), leading to a mean heritability around 0.25, a minimum heritability of 0.02, and a maximum heritability of 0.49. Therefore, we simulated the IDP-heritability between 0.02 and 0.25. For the phenotype heritability (corresponding to the SNVs under an IDP), relatively low values were selected (0.025, 0.05 and 0.1); this is because of the assumption that the overall phenotype heritability is contributed to by multiple IDPs and other factors, leading to a relatively lower heritability contributed by the genetic variants underlying a single IDP.

Under both pleiotropy and causality scenarios, we observed that the heritability of IDP or phenotype is relevant to the powers of all methods. Although the power of IMAS dropped a bit with an increasing number of causal SNVs when the heritability is relatively low, the overall power ranges from 0.88 to 1.0 under pleiotropy architecture and 0.83 to 1.0 under causality architecture (**Figure 5**).

**Figure 5.**
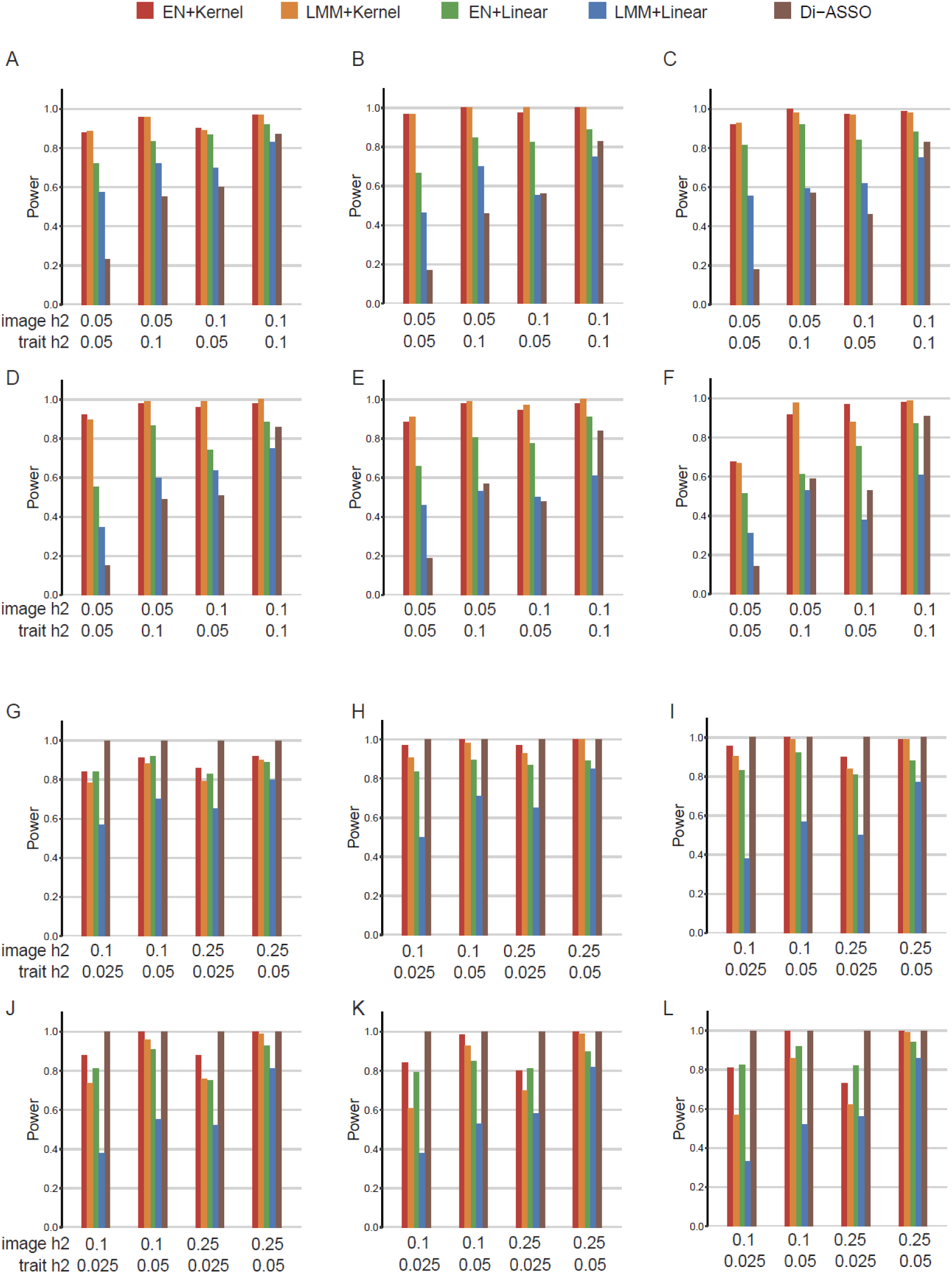
Power comparison of protocols in simulated pleiotropy and causality scenarios. Power is indicated on the y-axis. All panels are results under an additive genetic architecture, with differing image heritability and local trait heritability denoted below each panel. The total number of contributing genetic variants is 5, 10, 20, 50, 75 and 100 in panels AG, BH, CI, DJ, EK, FL, respectively. A-F. The pleiotropic scenario, which simulates independent associations from genotype to phenotype and expressions. Di-ASSO is short for the Direct-association model. G-L. The causality scenario, which simulates dependence of phenotype on genotype via gene expression.

Under the pleiotropy scenario, it’s evident that the power of Di-ASSO is substantially suboptimal compared with IMAS protocols (**Figure 5A-F**). An explanation is that when the effect of genetic variants on phenotypes is not causal through the IDP, the non-genetic effects within the real IDP add noises to the association test. In contrast, the IMAS method captures the genetic component of IDPs, leading to a higher power. Under the casualty scenario, Di-ASSO generally outperforms IMAS (**Figure 5G-L**). However, the power of IMAS is still comparable with Di-ASSO, especially when the IDP and phenotype heritability are relatively high. Notably, the power of Di-ASSO is all 1.0 based on the parameterization under our hypotheses. In practice, we expect that brain images may be a biomarker for disease progression and not necessarily the cause of the disease. As such, we expect pleiotropy scenarios may be more prevalent. This result has significant practical implication given that brain imaging, e.g., MRI, can be prohibitively expensive. In practice, IMAS can be used as a replacement for the standard test that requires images to be assessed, or at least a preliminary statistical analysis before conducting imaging.

## DISCUSSION

IMAS is a new method that leverages existing imaging data (such as IDPs in the UK Biobank) to mediate genotype-phenotype association studies. With empirical data analysis, we showed that IMAS can identify IDPs relevant to the focal disorders, despite the IDPs not being assessed in the GWAS dataset. Additionally, the underlying SNV-set selected by corresponding IDPs are enriched in neuropsychiatric pathways and the SNVs that are eQTLs indeed regulate functionally important genes. Those images that are identified by our model indicate the possible regions implicated for the four diseases and may help distinguish different diseases with similar symptoms. The corresponding SNVs provide preliminary candidates for further functional studies towards the mechanism of neuropsychiatric disorders.

Although the cerebellum has long been believed to be critical for disorders of motor functioning^76^, it is increasingly recognized as one of the most important brain^24, 77, 78^ regions playing a crucial role in psychopathology of non-motor disorders as well. Similar to our findings, Kim *et al.* suggested that the disruption of white matter connectivity in the cerebellum has strong association with cognitive impairments in patients with schizophrenia^79^; and Mwangi *et al.* indicated that white matter in the cerebellum is predictive of the staging or neuroprogression of bipolar disorder^80^. Many irregularities in grey matter volume have been shown to be associated with bipolar disorder compared to healthy controls^81, 82^. Romer et al. reported that shared risk for a wide range of mental disorders is strongly associated with white matter abnormalities of the pons, which is in turn positively correlated with cerebellar grey matter volume. Taken together, these findings indicate that the cerebellum is involved in psychopathology of many mental disorders.

As demonstrated by simulations, when the focus is the genetic basis, “borrowed” images from an existing dataset such as the UK Biobank can work well (or even better in many scenarios) compared to having actual images in one’s dataset. This will bring substantial cost savings in practice. Additionally, researchers and clinicians can use IMAS to conduct a first round of analysis before deciding whether imaging is necessary, and which imaging techniques and which patients may need to be assessed with neuroimaging based on the *in-silico* results. In this work IMAS and standard GWAS (i.e., LMM) outcomes are compared with the real data analysis without further verification in simulations – this is because the power of IMAS and LMM largely depends on the level of genetic basis of IDP, which is somewhat arbitrary to specify in simulations.

IMAS is partly inspired by transcriptome-wide association study (TWAS)^10^ in the sense they both utilized endophenotype data, which is gene expression in TWAS and images in IMAS. There are three conceptual advantages of IMAS over TWAS. First, TWAS relies on the concept of genetically regulated expression (GReX) that serves as both the target of genotype-based prediction of expressions and the statistic of aggregating SNVs for the next step’s association mapping^17, 18^. In contrast, IMAS splits these two essentially independent steps into image-directed SNV selection and kernel-based SNV aggregation.

Second, TWAS is gene-based, therefore naturally it is straightforward to predict the expression levels mainly using local cis-SNVs, and it is unclear whether extending this concept to the whole-genome without further *a priori* knowledge may work well. In this work, by analyzing real data using kernel-based SNV aggregation in the association test, it has been proven that extending the concept works. Third, an inherent problem of TWAS lies in that the expression heritability is generally low^10^, therefore the power may be only marginally higher or even lower than GWAS in some scenarios^19^. However, the high heritability of IDPs (**Supplementary Figure 10**) over expression heritability allows a significantly higher power of IMAS over GWAS, as demonstrated in our real data analysis (**Supplementary Table 6-7**).

In 2017, Xu et al. published a tool to utilize imaging endophenotype to identify genes contributed to brain disorders, which they termed Imaging-Wide Association Study, or IWAS^83^. IWAS mimics the practice of TWAS in that it still predicts the contribution of cis-SNVs in a single gene to image(s) and then associates the gene to the disease. They have not touched the topic of selecting SNVs from the whole-genome, nor did they exhaustively analyze the advantage of higher heritability of IDPs as well as the striking advantage of “borrowing” images over the scenario of assess the real image.

Following the standard TWAS pipeline, BrainXcan^84^ has been proposed to analyze behavioral and psychiatric traits using large scale genetic and imaging data. The BrainXcan pipeline is similar to our IMAS protocol (1.a) + (2.a), except for its support to summary statistics whereas IMAS assumes subject level genotypes. In our simulations, this protocol is the least powerful among the four protocols (**Figure 5**). By applying BrainXcan^84^ to the four brain disorders, we were not able to identify any significant IDPs (**Supplementary Notes IV**).

That said, BrainXcan indeed enjoys the advantage of supporting summary statistics. IMAS can also be extended to summary statistics based on the idea of metaSKAT^85^. However, in the initial GWAS, researchers in general generate summary statistics using Z-score or other means without involvement of kernel, whereas metaSKAT assumes that the summary statistics in individual GWAS cohorts were generated by the kernel method (SKAT). We have not derived the model to carry out meta-kernel analysis using standard summary statistics. As such, before the community is convinced to provide summary statistics by kernel methods for the first line GWAS (which in our opinion is beneficial), it is unlikely that IMAS could be extended to summary statistics, which is a limitation.

Future work could include integration of more specific image analyses with genetic mapping and developing methods for integrating both images and gene expression into the same statistical framework.

## Acknowledgement

This work is supported by an Alberta Innovates LevMax-Health Program Bridge Funds (222300769 Q.L. & P.A.), a New Frontiers in Research Fund grant (NFRFE-2018-00748, Q.L.), and an HBI Pilot grant (Q.L. & P.A.). J.H. is supported by a CSC scholarship, A.L. is supported by Heritage Youth Researcher Summer Program. P.A. also receives support from the Alberta Innovates Translational Health Chair in Child and Youth Mental Health. The computational infrastructure is supported by a Canada Foundation for Innovation JELF grant (36605).

## Author contributions

(Q.L. = Quan Long, Q.L.2 = Qing Li) Conceived the study: P.A. and Q.L.; Developed the software: J.H.; Conducted numerical simulations: J.H., W.Z.; Analyzed real data: J.H., Q.L.2, D.E.; Provided biological interpretation: L.A., H.Z., A.L. and P.A.; Provided comments: B.C. and M.E.M. Supervised the study: P.A. and Q.L. Wrote the paper: J.H., L.A., P.A. and Q.L. with the input from all coauthors.

## Web resources

IMAS, https://github.com/theLongLab/IMAS

mkTWAS, https://github.com/theLongLab/mkTWAS

kTWAS, https://github.com/theLongLab/kTWAS

PrediXcan, https://github.com/hakyim/PrediXcan

BrainXcan, https://github.com/hakyimlab/brainxcan

JAWAMIX5, https://github.com/theLongLab/Jawamix5

PLINK, https://zzz.bwh.harvard.edu/plink/

GCTA, https://yanglab.westlake.edu.cn/software/gcta/#Download

cS2G mapping: https://alkesgroup.broadinstitute.org/cS2G/cS2G_UKBB/

Qiagen Ingenuity Pathway Analysis (IPA), https://digitalinsights.qiagen.com/products-overview/discovery-insights-portfolio/analysis-and-visualization/qiagen-ipa/

UK BIOBANK, https://www.ukbiobank.ac.uk/

1000 Genomes Project, https://www.internationalgenome.org/

GTEx, https://gtexportal.org/ and https://www.ncbi.nlm.nih.gov/projects/gap/cgi-bin/study.cgi?study_id=phs000424.v8.p2

Genome-Wide Association Study of Schizophrenia, https://www.ncbi.nlm.nih.gov/projects/gap/cgi-bin/study.cgi?study_id=phs000021.v3.p2

Major Depression: Stage 1 Genome-wide Association in Population-Based Samples, https://www.ncbi.nlm.nih.gov/projects/gap/cgi-bin/study.cgi?study_id=phs000020.v2.p1

Whole Genome Association Study of Bipolar Disorder, https://www.ncbi.nlm.nih.gov/projects/gap/cgi-bin/study.cgi?study_id=phs000017.v3.p1

MSSNG, https://research.mss.ng/

MEDLINE database, https://www.nlm.nih.gov/medline/medline_overview.html

## ONLINE METHODS

### The IMAS model and implementation

The IMAS model includes two steps. First, IMAS selects genetic variants (and their weights) using the IDP cohort that contains genotype and the image features. Second, IMAS aggregates the selected (weighted) variants to associate them to the disease phenotype. For each step, IMAS provides two alternative methods (1.a), (1.b) and (2.a), (2.b), potentially four configurations. However, the recommended default protocol is (1.b) + (2.b), i.e., linear mixed model plus kernel method.

***Step 1:*** For feature selection, we implemented two models: regularized regression (Elastic Net^10^) and linear mixed model (EMMAX^40, 41^).

<u>(1.a) Elastic Net:</u>

An additive model is used to fit a linear combination of genetic variants:

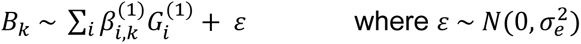

where superscript (1) indicates the Step 1 dataset, *B* is the brain image matrix, *B_k_* is the *k*-th IDP, *G*(1) is the genotype matrix, *G_i_*^(1)^ is the number of reference alleles of *i*-th variant (=0, 1, or 2), *β_i,k_*^(1)^ is the effect size of *i*-th variant for *k*-th IDP, and ɛ is the residual with *σ_e_*^2^ as its variance. Using a mixture of *L*_1_ and *L*_2_ regularization, we solve the (*β_i,k_*^(1)^) by Elastic Net, a regularized regression model. The variants with non-zero coefficients will be selected.

<u>(1.b) Linear Mixed Model:</u>

In contrast to the above method using a linear combination of variants, feature selection is carried out by assessing marginal effect of each SNV individually. This is achieved by fitting a linear mixed model:

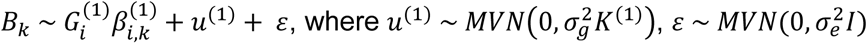

Where *B_k_*, *G*^(1)^, *β_i,k_*^(1)^ are the same as defined in (1.a); *u*^(1)^ is the random effect term with *var*(*u*^(1)^) = *σ_g_*^2^*K*^(1)^, where *K*^(1)^ is the genetic relationship matrix (GRM), which is calculated by *G*^(1)*T*^*G*^(1)^/*n*, where *n* is the number of SNVs, *ε* is an the residual effect such that *var*(*ε*) = *σ_e_*^2^*I*. *MVN* denotes multivariate normal distribution.

This model is equivalent to EMMAX^40^ and in IMAS our out-of-core implementation^41^ is utilized. The SNVs with nominal *P-value* lower than a user-prespecified cutoff (default =0.01) will be selected.

***Step 2:*** The method for aggregation may be multiple linear regression (as in the original TWAS protocols^10^) or a kernel machine^37^ which is more robust to noise, as showed by previously^17, 18^.

<u>(2.a) Linear Combination:</u>

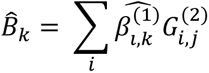

where superscript (2) indicates the Step 2 dataset, *B̂_k_* is the estimated IDP values for *k*-th IDP in the GWAS cohort. 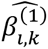 is the coefficient for *i*-th variant and *k*-th IDP derived from Step 1. *G*^(2)^ is the genotype in the GWAS cohort, in which *G_i,j_*^(2)^ is the variant of the *i*-th selected variants of the *j*-th individuals. *B̂*_*k*_ will then be tested for association with the phenotype in the GWAS cohort.

<u>(2.b) Weighted Kernel:</u>

IMAS forms a weighted kernel using the selected SNVs and their estimated coefficients:

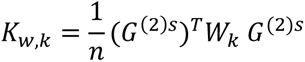

Where *G*^(2)*s*^ is standardized genotype matrix using *G*^(2)^ (by rescaling each SNV’s mean to be 0 and variance to be 1), 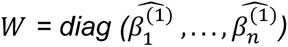, where *n* is the number of selected variants. The subscript *k* denotes the *k*-th IDP.

This weighted kernel reflects the similarity of subjects in terms of SNVs that are functionally relevance to images, which will be used for association test as implemented in the sequence kernel association test (SKAT)^37, 86^:

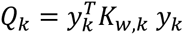

where *y_k_* is the vector of phenotype values, and *K_w,k_* is calculated above. This score *Q_k_* follows a mixture of chi-squared distribution^37, 86^.

We recommend allowing a relatively large number of potential SNVs (e.g., SNVs with nominal p-values < 0.01 before multiple-test correction) to be aggregated in the second SKAT. This is because our previous work in TWAS revealed that the performance of SKAT favors a large number of weakly associated variants over a small number of highly significant variants^17^, possibly due to the advantage of kernel methods being more robust to noise.

### Procedure of Simulations

#### Genotype data

Based on the real UK Biobank genotype (containing 33,553 individuals with 367,986 SNVs), we simulate brain images in the reference dataset (for IMAS Step 1 analysis). Based on the 1000 Genomes Project^87^ genotype (containing 2,548 subjects with 4,422,985 SNVs), we simulate phenotype in the GWAS dataset (for IMAS Step 2 analysis).

As we also compare with the IMAS power to the hypothetical protocol where brain images are available in the GWAS dataset, brain images are also simulated using the 1000 Genomes Project genotype. We assume that the brain image data is quantitative, same as the format of UK Biobank IDPs. In all cases, we simulate a genetic component first and then incorporate it to the IDP or phenotypes using prespecified values of genetic component (i.e., heritability).

#### Genetic models underlying IDP and phenotype

Two typical genetic architectures are considered: causality, where genotypes alter phenotype via the image (Supplementary Figure 11A) and pleiotropy, where genotypes contribute to phenotype and image independently (**Supplementary Figure 11B**).

Under both scenarios, we simulate IDPs and phenotypes with an additive genetic architecture, in which phenotypes and IDPs are associated with the sum of genetic effects. We first randomly selected a prespecified number of SNVs (5, 10, 20, 50, 75, 100) with minor allele frequency (MAF) higher than 1% as causal variants, and then simulate the genetic component of IDPs or phenotypes below:

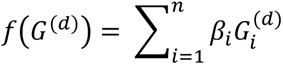

where *G* denotes the genotype matrix, The effect sizes *β_i_* are drawn from the standard normal distribution *N*(0,1). Superscript *d* may be 1 or 2 for reference (UK Biobank) and GWAS datasets (1000 Genomes Project) respectively. The IDPs in reference and GWAS datasets thus are generated by:

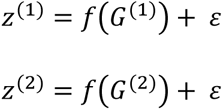

where *ε*^(*IDP*1)^ and *ε*^(*IDP*2)^specify the levels of genetic component by adjusting their variance (will be detailed below).

With a causality model, the phenotype is decided by IDPs:

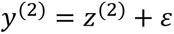

Whereas with a pleiotropy model, the phenotype is decided by genetics directly using the same formula for IDPs (except that the variance component is rescaled by image heritability instead of trait heritability):

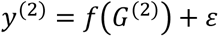

The genetic component contributing to IDPs or phenotypes is prespecified as a value between 0 and 1. Under both models, the variance components are rescaled by image heritability and phenotype heritability to ensure the relevant heritability indeed matches the prespecified parameters.

#### Generating residuals to warrant the heritability being specified values

In the above simulations of IDP and phenotypes, the residual *ε* serves as the role to adjust genetic components based on prespecified values (i.e., heritability) denoted as *h*^2^. To achieve that, we first generate the genetic component based on related formulas and real genotypes and calculate its variance of the genetic component as *σ_g_*^2^. We then solve *σ_e_*^2^ in equation 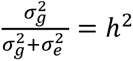. We then sample *ε* ∼ *N* (*O*, *σ_e_*^2^) to determine values of residuals, ensuring the heritability to be exactly *h*^2^.

#### Type-I error estimation & power calculations

We generated random phenotypes using genotype data from the 1,000 Genomes Project to form the empirical null distribution of the p-values. Then the type-I error for the IMAS model was estimated using the top 5% cutoff for the most significant p-values.

For each of the genetic architectures and their associated parameters, we simulated 1,000 datasets each, in which causal variants are randomly selected. The power was then calculated as the proportion of each protocol’s success in identifying the IDPs in each dataset, where the success is defined as a Bonferroni-corrected p-value that is lower than a predetermined critical value (0.05).

#### Direct association analysis in the hypothetical protocol where IDPs are available in the GWAS dataset

A goal in our simulation is to compare the power of IMAS that borrow IDPs from reference dataset to a naïve method that directly associate IPDs to phenotype (assuming IDPs are available in the GWAS). Towards this line, a simple linear regression is used to associate IDPs with phenotype:

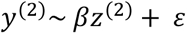

where *y*^(2)^ is the simulated phenotype and *z*^(2)^ is the simulated image phenotype in the GWAS dataset (based on the 1,000 Genome Project genotypes).

### Real Data Analysis

#### Data source and quality control

The UK Biobank IDPs are downloaded from their website under the application number #ukb45132 and the IRB #REB19-1121. Initially, we collected 34,568 individuals in UK Biobank with 885 IDPs and non-missing values for other covariates such as genetic PCs, sex, and age at recruitment. First, each IDP’s data vector had outliers removed (set to missing, with outliers determined by being greater than six times the median absolute deviation from the median)^88^. Then, we discarded individuals for whom 50 or more IDPs were missing^88^. Each IDP’s data vector was normalized, resulting in it being Gaussian distributed, with mean = 0 and s.d. = 1. Lastly, we regressed out covariates, including the top 10 genetic principal components supplied by UKB (UK Biobank DataField 22009), age, sex, squared age, age × sex, squared age × sex, head size (UK Biobank DataField 25000), head motion (UK Biobank DataField 25741), and scanner positions (UK Biobank DataField 25756, 25757, 25758, 25759)^88, 89^. Eventually, we collected 33,553 individuals in UK Biobank for 885 IDPs.

We run IMAS with default parameters on four different brain diseases datasets, i.e., SCZ^20^, MDD^21^, BPD^22^ and ASD^23^, downloaded from dbGaP web portal (**Web resources**) and the MSSNG database (**Web resources**) ^23^. All genotyping datasets are quality controlled using PLINK^90^. The QC steps involved: excluding markers with missingness rate > 0.1, minor allele frequency < 0.05, high deviations from Hardy-Weinberg equilibrium>10^-6^ and removing samples with missingness rate > 0.1. The significant IDPs are defined as genome-wide significant after Bonferroni correction of *P*-value < 0.05. The multiple-test correction considers the number of IDPs. Eventually, we collected 2,548 individuals with 728,620 SNVs for SCZ, 3,644 individuals with 450,236 SNVs for MDD, 1,028 individuals with 844,975 SNVs for BPD, and 7,068 individuals with 5,140,328 SNVs for ASD after quality control.

When aligning two datasets’ genotype coordinates, i.e., the reference dataset (UK Biobank) and GWAS dataset (the 4 disorder cohorts), Liftover^91^ was utilized to convert all coordinates to HG38. The locations that are in overlap are used for the analyses.

#### Mapping SNVs to genes and pathway enrichment analysis

To validate the functional relevance of the identified IDPs with focal diseases, we first conducted literature research for evidence from other studies. Then, we carried out GO enrichment and KEGG pathway analysis to confirm our findings. We first collected top SNVs (P < 0.01) for each significant IDP (that is associated with the focal disorder), then we mapped those SNVs to genes according to SNV-to-gene linking reported by Steven Gazal, et al^92^. After mapping SNVs to (a set of) genes, we conducted GO enrichment^93^ and KEGG pathway analysis^94^. The GO enrichment is carried out from three different aspects: ‘Biological Process’ (BP), ‘Cellular Component’ (CC) or ‘Molecular Function’ (MF)^93^. KEGG is used to systematically analyze gene functions, which links gene information with high-level functional information^95^. The GO enrichment and KEGG pathway analyses were conducted using the ‘clusterProfiler’ R package^96^. The enriched GO terms and KEGG pathways with a p < 0.05 were considered disease-related biologic processes or signaling pathways.

#### Standard GWAS analysis for the four disorders

The standard GWAS analyses of four disorders are conducted using standard linear mixed model (LMM) ^40, 41^. Mathematically, it is the same as the feature selection part (1.b) in IMAS, expect for that we used phenotype instead of IDP as the dependent variable *y* :on the left-hand side of the regression.

#### eQTL analysis

To link the IDPs to gene regulatory mechanisms, we conduct eQTL analysis using eQTL files of brain tissues downloaded from the GTEx Consortium^97^. For each significant IDP, we first mapped selected SNVs associated with this IDP to expression quantitative trait loci (eQTL), SNVs whose genetic variation explains a fraction of the variance of gene expression level. The criterion of the mapping is that the IDP-associated SNV and the eQTL must share identical location. Then, for the SNVs that we can successfully map to an eQTL, we rank the genes being regulated using the level of significance in LMM-based feature selection (step (1.b) of IMAS). These regulated genes were then deemed as genes explaining the genetic basis of brain disorders medicated by IDPs via the expression regulatory mechanism. To verify such mechanisms, we searched for literature proofs of top five regulated genes (**Supplementary Notes III**).

#### Cross-disorder co-expression matrix of cerebellum-related genes

Based on the above eQTL analysis, we first prioritized the top 20 eQTL genes with the most significant p-values for each disorder. Then for each pair of disorders (e.g., MDD and SCZ, as shown in **Figure 4B**), using the GTEx gene expression data (in the cerebellum tissue), we calculated the Pearson’s correlations between expressions of every pair of genes (in total 20×19/2 = 190 pairs). This forms the cross-disorder co-expression matrices for all pair of disorders, supporting the heatmaps in **Figure 4B**; **Supplementary Figures 5-9**.

#### Gene-to-gene connection network

The gene-to-gene connection network was constructed by searching literature and databases based towards the genes identified in the co-expression matrices. Multiple resources such as MEDLINE database^98, 99^ and KEGG pathway database^94^ were used. We included the gene-to-gene connections that are associated with neurological traits or play a role in molecular, metabolomic and immunological cellular functions, and excluded those connections that have only been reported in the cancer cell lines. The network was generated via Qiagen Ingenuity Pathway Analysis (IPA) software^100^.

