## Supplementary materials for "A statistical method for image-mediated association studies discovers genes and pathways associated with four brain disorders"

### Supplementary Notes

#### Supplementary Notes I: Annotations of IMAS-discovered IDPs.

***Schizophrenia*.** we observed three significant IDPs, i.e. Mean MO in fornix cres+stria terminalis on FA skeleton (IDP#=25190-2.0, $P=9.82\times10^{-6}$ ), Mean L3 in inferior cerebellar peduncle on FA skeleton (IDP#=25306-2.0, $P=3.69\times10^{-5}$ ), and Mean FA in fornix cres+stria terminalis on FA skeleton (IDP# = 25094-2.0, nominal $P=5.44\times10^{-5}$) (**Figure 3A**).

[IDP-25190-2.0, Mean MO in fornix cres+stria terminalis on FA skeleton] The fornix+stria terminalis of the limbic system plays a large role in the regulation of behavior and emotion in an individual. However, there have been less studies on the connection between the fornix+stria terminalis and SCZ. Lee et al. suggests that there exists a correlation between the fornix+stria terminalis and SCZ. It was discovered that FA values of the right fornix cres / stria terminalis in the chronic SCZ group were significantly lower than those of the healthy control group[1].

[IDP-25306-2.0, Mean L3 in inferior cerebellar peduncle on FA skeleton] The cerebellum communicates with the other central nervous system through the three major white matter (WM) bundles: superior (SCP), middle (MCP), and inferior (ICP) cerebellar peduncles[2]. These cerebellar peduncles have an essential role in integrating the cerebellum into distributed neural systems to modulate and optimize cognitive, affective, and sensorimotor activities through trial-and-error learning[2, 3].

[IDP-25094-2.0, Mean FA in fornix cres+stria terminalis on FA skeleton] A study has reported reduced FA values in the fornix and stria terminalis in chronic SCZ when compared to first episode SCZ and healthy controls, which suggested that fornix and stria terminalis play an important role in pathophysiology of SCZ[1]. Another study also showed that patients with schizo-obsessive comorbidity, a subgroup of SCZ, had decreased FA and increased radial diffusivity in the left crescent of the fornix and stria terminalis compared with healthy controls[4].

***Major depression disorder*.** Eight significant IDPs were listed in Table 1 (**Figure 3D**). In the below context, we annotated the top four most significant IDPs that are associated with MDD.

[IDP-25152-2.0: Mean MO in middle cerebellar peduncle on FA skeleton] The majority of results from diffusion imaging studies showed a weaker structural connectivity in participants with ASD, as indicated by decreased FA both in the superior cerebellar peduncles[5-7] and in intracerebellar circuitries[8]. However, findings in the middle cerebellar peduncles did not yield consistent evidence. Shukla et al[9]. revealed reduced values of FA in adolescents with ASD; conversely, Sivaswamy et al[10]. found increased values of FA in the right middle cerebellar peduncle, although within a reverse asymmetry pattern in FA of the middle and inferior cerebellar peduncles[11].

[IDP-25905-2.0: Volume of grey matter in Crus II Cerebellum] Reduced grey matter volume represents a hallmark of MDD neuropathology, typified by a wide-ranging distribution of structural alteration. Compared MDD patients with healthy controls, GM volume alterations have been associated with cognitive deficits[12], sustained attention reduction[13], executive processing[14], anxiety disorders[15], and emotional dysregulation[16]. However, there are controversial ideas on the key regions of GM structural abnormality. Volumetric reductions have been reported in different brain regions, including cerebellum[14]. Mujeeb et al. showed that depressed adolescents had lower gray matter volume in bilateral cerebellum (uvula and tonsil) than controls[17], an earlier study reported that decreased GM volume in the left cerebellum in patients with first-episode depression[18] and Stuart et al. also found widespread reductions in GM volume in MDD participants, including cerebellum[19]. Although MDD-related symptoms and GM reduced regions are highly heterogeneous, GM alteration appears to be a constant consequence.

[IDP-25862-2.0: Volume of grey matter in Frontal Operculum Cortex] The frontal operculum rostral to the ascending ramus of the lateral fissure is associated with the prefrontal association cortex and plays a role in thought, cognition, and planning behavior. Studies indicate that the prefrontal cortex has emerged as one of the regions most consistently impaired in major depressive disorder (MDD)[20]. Cotter et al. indicated that area 9 (see figure) of the dorsolateral prefrontal cortex in individuals with major depressive disorder have reduced neuronal size and glial cell density[21]. Furthermore, consistent with our findings, researchers have discovered findings of gray matter abnormalities in frontolimbic areas in depressed adults. This suggests the presence of these structural changes at the onset of depressive illness[17].

[IDP-25703-2.0: Weighted-mean OD in tract uncinate fasciculus]: Many studies have shown that the alterations in white matter (WM) microstructure are involved in the pathophysiology of MDD[22]. Uncinate fasciculus, known as white matter fiber tract, connecting brain structures, has been one of most well-studied regions associated with depression[23]. Studies have shown that patients with MDD had lower FA in the tract connecting subgenual anterior cingulate cortex (ACC) to amygdala in the right hemisphere, and the tract is likely encompassed by the uncinate fasciculus, whose fibers connect the medial temporal cortex with the orbitofrontal cortex, including the subgenual ACC[24]. The subgenual ACC plays an important role in regulating emotion, and degeneration in this area correlates with depressed mood and anhedonia[25].

**Autism spectrum disorder.** Fourteen significant IDPs were listed in Table 1 (**Figure 3G**). In the below context, we annotated the top four most significant IDPs that are associated with ASD.

[IDP-25869-2.0, Volume of grey matter in Planum Polare] The structure of the planum temporale is often disturbed in disorders with associated communication problems, particularly in auditory and language processing areas in the brain. The volume of gray matter in the planum temporale was reduced in the ASD group, which suggest an early neurodevelopmental disturbance in ASD[26-28].

[IDP-25096-2.0, Mean FA in superior longitudinal fasciculus on FA skeleton & 25288-2.0, Mean L2 in superior longitudinal fasciculus on FA skeleton] One of the most reported neural features of ASD is the alteration of multiple long-range white matter fiber tracts, as indexed by reduced fractional anisotropy (FA)[29]. Studies have found that reduced FA of superior longitudinal fasciculus (SLF) had negative correlations with social interaction, which contributes to both core and associated symptoms of ASD. Alterations in the SLF has also been reported to be associated with language impairment in ASD[30].

[IDP-25196-2.0, Mean MO in uncinate fasciculus on FA skeleton] The uncinate fasciculus (UF) is a long-range white matter tract that connects limbic regions in the temporal lobe to the frontal lobe. Abnormality of the UF have been reported in numerous studies of individuals with developmental and psychiatric disorders, including ASD[31]. Many studies suggested that abnormalities of the UF are present in individuals with ASD that could contribute to the characteristic deficits seen in social and communications deficits, difficulties regulating emotions as well as cognitive functioning[32-40].

[IDP-25288-2.0, Mean L2 in superior longitudinal fasciculus on FA skeleton] Many studies have found that impairment of neural connectivity is associated with social and communications deficits and cognitive impairments in ASD[30, 41]. A Study has shown that decreased FA of the inferior longitudinal fasciculus (ILF) and superior longitudinal fasciculus (SLF) had negative correlations with scores of social interactions[41]. Language impairments are observed in a subset of individuals with ASD. Abnormal DTI parameters (specifically significantly elevated MD values in ASD) of the superior longitudinal fasciculus appear to be associated with language impairment in ASD[30].

***Bipolar Disorder.*** The only significant IDP found for BPD was the Volume of grey matter in Vermis Crus II Cerebellum (IDP# = 25904-2.0; $P=2.20\times10^{-5}$). We also identified the same IDP for MDD, which explains the similarity between these two disorders (**Figure 3J**).

A growing number of papers have shown that the cerebellum modulates affect and cognition in addition to motor functions, which contributes substantially to the pathophysiology of mood and psychotic disorders, such as BPD[42, 43]. Patients with BPD showed a decline in cerebellar grey matter density at an accelerated rate compared with healthy control subjects, and this tissue loss is associated with deterioration in cognitive function and illness course[44]. However, similar to what we have found for MDD, the findings on grey-matter volume alterations in BPD are also heterogeneous. Besides cerebellum, some studies reported the grey-matter abnormalities in the prefrontal cortex and insula[45], which may be due to the differences in specific demographic and clinical features of each patient[45].

**Supplementary Notes II: Pathway enrichment analysis of IMAS-identified SNVs.**

***Schizophrenia*.**

[IDP-25094-2.0] Focal adhesion was one of the leading pathways in KEGG analysis. Focal adhesions have been shown to play a role in regulating cellular behaviors such as cell migration, stabilizing the cell during this process, and creating combinatorial signaling complexes as well as mediating integrin function[46]. Focal adhesion kinases and their downstream signaling pathways also have been shown to play a role in cell migration and angiogenesis[47]. Studies on the olfactory neutrospheres-derived cells of SCZ patients revealed altered motility and focal adhesion dynamics which support evidence of dysregulation expression of genes in focal adhesion kinase signaling in SCZ patients[48]. The significantly enriched GO terms for biological processes are the regulation of cell-cell adhesion, cell-substrate adhesion, ameboidal-type cell migration, neuron migration, and neural crest cell migration. The significantly enriched GO term for cellular components is focal adhesion.

[IDP-25306-2.0] PI3K-Akt signaling pathway was one of the leading pathways in KEGG analysis. The PI3K-Akt signaling pathway plays an indispensable role in the cell for proliferation and apoptosis. This pathway also plays a role in cell migration and vesicle transport[49]. Evidence from observing decreased Akt-1 protein and its kinase in white blood cells and brain tissue of schizophrenic patients has indicated the involvement of the Akt signaling pathway in the pathogenesis of this disorder. Interactions between the Akt signaling pathway and genes responsible for neurodevelopment, synapse formation, and synaptic plasticity explain the effect on the pathogenesis of SCZ[50]. Other pathways, including Focal adhesion, have also been reported to be involved in the pathogenesis of SCZ. The focal adhesion pathway was another leading pathway in KEGG analysis for IDP-25306-2.0. This pathway was the leading pathway identified in IDP-25094-2.0, IDP-25284-2.0, and other IDPs for SCZ. The significantly enriched GO terms for biological processes are neuron projection guidance, negative regulation of neurogenesis, negative regulation of neuron differentiation, neuroblast proliferation, negative regulation of excitatory postsynaptic potential, and neural precursor cell proliferation.

[IDP-25190-2.0] The rap1 signaling pathway was one of the leading pathways in KEGG analysis. The Rap1 signaling pathway plays an important role in the control of platelet activation and neuronal plasticity, intersecting with Ca2+ signaling pathways to do so. In the central nervous system, Rap1 is involved in numerous Ca2+-dependent processes which include synaptic plasticity, cotico-amygdala plasticity, and long-term potentiation and gene transcription[51]. Mental disorders causing learning disability have been associated with genetic mutations of molecules involved in Rap signaling demonstrating the role of Rap signaling in human learning and memory[52]. Impairment in the capacity of synaptic plasticity has been observed in cases of both hypo- and hyperactivation of Rap signaling[52]. Additionally, various genetic defects that up- or down-regulate Rap signaling have been linked to mental disorders such as SCZ which are associated with learning disabilities[52]. Other pathways, including focal adhesion, have also been reported to be involved in the pathogenesis of SCZ. The focal adhesion pathway was another leading pathway in KEGG analysis for IDP-25306-2.0. This pathway was the leading pathway identified in IDP-25094-2.0, IDP-25190-2.0, IDP-25306-2.0, and IDP-25284-2.0 other IDPs for SCZ The significantly enriched GO terms for biological processes are the learning or memory, and regulation of transcription involved in cell fate commitment.

***Major depression disorder*.**

Below, we annotated the top four most significant IDPs that are associated with MDD.

[IDP-25152-2.0] Axon guidance was one of the leading pathways in KEGG analysis. The Axon guidance signaling pathway plays an important role in allowing axons to form synaptic connections. The pathway plays a role in growth cones and axons, rearranging the local cytoskeleton and plasma membrane[53]. A recent study has found that axon guidance signaling was a significantly altered pathway in Major depressive disorder patients[54]. Axon guidance genes and proteins are involved in the pathogenic mechanisms of neuropsychiatric disorder as they play an important role in controlling the migration and synapse plasticity of neuronal cells[55, 56]. It has been found that Netrin-1 and its receptor DCC are elevated in the cortex of MDD patients, these establish synaptic connections by acting as axon guidance cues[57]. The significantly enriched GO terms for biological processes are axon guidance, axon extension involved in axon guidance, axonogenesis, regulation of axon extension involved in axon guidance, and regulation of axon guidance.

[IDP-25703-2.0] The calcium signaling pathway was one of the leading pathways in KEGG analysis. Calcium signaling pathways are important in controlling a diverse array of processes such as metabolism, secretion, fertilization, proliferation, and smooth muscle contraction. The signaling pathway can induce changes in the generation and function of the $Ca2+$ signal. Disturbed brain development, neuroplasticity, and chronobiology are all factors associated with BPD, one of the causes of these characteristics is calcium signaling pathways. Whole genome sequencing has supported the role of calcium in BPD with this disorder being associated with calcium dysregulation and apoptosis. One hypothesis for BPD is the calcium and mitochondrial dysfunction hypothesis where mitochondrial dysregulation of $Ca2+$ leading to symptoms of BD are caused by nuclear gene mutations that create mtDNA polymorphisms/mutations or mtRNA deletions[58]. The significantly enriched GO terms for biological processes are the regulation of synaptic plasticity.

[IDP-25905-2.0] cAMP signaling pathway was one of the leading pathways in KEGG analysis. Transcription factor cAMP response element-binding protein (CREB) plays a role in cognition, underlying learning, and memory. Additionally, CREB mediates gene expression which is required for long-term memory and synaptic plasticity[59]. Examining the antidepressant activity of Huang-lian Jie-du Decoction (HLJDD) on chronic unpredictable mild stress-induced depressive mice and was found to be implicated in the inhibition of depressive symptoms through modulating the Camp signaling pathway among other pathways[60]. Research has found that the inhibition of the ERK pathway in the prefrontal cortex and hippocampus causes depression-like behavior with the cAMP response element-binding protein (CREB) being vulnerable to depression. CREB is reduced in activity in depressed animals[61]. The significantly enriched GO terms for biological processes are positive regulation of neuron differentiation, positive regulation of neurogenesis, developmental cell growth, axonogenesis, and regulation of synaptic plasticity. The biological go terms of positive regulation of neurogenesis and regulation of synaptic plasticity relate to MDD as in this disorder there is a decrease in volume of the hippocampus. The two mechanisms proposed to explain this are that there is either an atrophy of mature neurons in the hippocampus lessening neuroplasticity or that new neurons are not formed because of a decrease in neurogenesis which leads to decrease of hippocampus volume in patients with MDD[62].

[IDP-25862-2.0] The Rap1 signaling pathway was one of the leading pathways in KEGG analysis. The Rap1 signaling pathway plays an important role in the control of platelet activation and neuronal plasticity, intersecting with Ca2+ signaling pathways to do so. In the Central nervous system Rap1 is involved in numerous Ca2+-dependent processes which include synaptic plasticity, cotico-amygdala plasticity, and long-term potentiation and gene transcription[51]. A study examining the levels of Rap1 protein in platelets of untreated euthymic and depressed patients with major unipolar depression found that Rap1 was significantly lower in untreated depressed patients when immunolabeling was conducted as compared to untreated euthymic patients and healthy subjects[63]. Another study looking at the rodent forebrain examined the Rap-1 guanine nucleotide exchange factor (MR-GEF) expression in individuals with major psychiatric disorders (SCZ, BPD, and MDD) as well as control individuals observing a positive correlation between the percentage of MR-GEF expressing neurons in individuals with BPD. This suggests changes in MR-GEF expression could influence neurotransmission[64]. The significantly enriched GO terms for biological processes are modulation of chemical synaptic transmission and regulation of synaptic plasticity. The biological GO terms that modulate chemical synaptic transmission relate to MDD as neurotransmission, is one of the best-known neurobiological correlates of this disorder[65].

***Bipolar Disorder.***

The only signal for BPD is IDP-25904-2.0 that has been also identified in MDD. 'Axon guidance' was one of the leading pathways in KEGG analysis (**Figure 3G**). Axon guidance signaling pathway plays an important role in rearranging the local cytoskeleton and plasma membrane in growth cones and axons, and it also controls gene expression via local translation and transcription[53]. Studies have shown proteins that regulate axon guidance were reduced in the regional hippocampus and entorhinal cortex of patients with BPD[66]. A more recent study has reported microRNAs that expressed in BPD-derived neurons were enriched in several cellular pathways, including axon guidance pathway[67]. Other pathways, including the leading pathway of PI3K-Akt signaling pathway have also been reported to be involved in the pathogenesis of BPD. The significantly enriched GO terms for biological processes are the regulation of axonogenesis, axon guidance, and regulation of membrane (**Figure 3H**).

***Autism spectrum disorder.***

Below, we annotated the top four most significant IDPs that are associated with ASD.

[IDP-25869-2.0] Focal adhesion was one of the leading pathways in KEGG analysis. Examining the molecular and physical mechanisms underlying cell migration, adhesion to substrate via specific focal adhesion points is considered an essential step in cell migration[68]. Focal adhesion kinase functions as an integrator to control cell motility and plays a key role in cell migration[69]. A study performing isobaric tags for relative and absolute quantitation analysis for control and autistic children's plasma found differentially expressed proteins as biomarkers for ASD. Among the biomarkers found most were involved in focal adhesion and other pathways[70]. Another study examined FAK (focal adhesion kinase) and found its expression was diminished in autistic lymphoblasts while adhesion and migration defects were restored with FAK overexpression. This is interesting considering that in the pathogenesis of ASD one of the major pathways is reduced cell migration and FAK plays an important role in many functions including neural migration[71]. The significantly enriched GO terms for biological processes are cerebral cortex radially oriented cell migration and cerebral cortex cell migration. The biological GO terms of cerebral cortex cell migration relate to ASD as one of the characteristics that is commonly observed in this disorder is deficiencies in neuronal migration as well as axon guidance[72].

[IDP-25096-2.0] The MAPK signaling pathway was one of the leading pathways in KEGG analysis. The MAPK pathway plays a developmental role in the development of the central nervous system and cerebral cortex. Literature links MAPK signaling to both Central nervous system development as well as many of the mature processes governing its functions. Another role of ERK/MAPK signaling in neurons is regulation of synaptic plasticity, an important mechanism where neural circuits can be modulated to change connections[73]. Studies have shown the role of ERK/MAPK signaling for the proper development of the nervous system and therefore ERK/MAPK's role in many neurodevelopmental disorders[73]. The ERK/MAPK pathway can be linked to ASD as it acts as a central hub that interacts with many of the genes and copy number variants that are implicated in ASD suggesting dysregulation of this pathway could contribute to the pathogenesis of ASD[73]. The significantly enriched GO terms for biological processes are axonogenesis, and axon guidance. The biological GO terms of axonogenesis relate to ASD as many of the ASD predisposition genes have been identified as being involved in the biological process of axonogenesis[74].

[IDP-25196-2.0] Axon guidance was one of the leading pathways in KEGG analysis. Axon guidance is a neurodevelopmental process that establishes neuron pathways and cortical circuits through the directing of growth cones[75]. The Axon guidance signaling pathway plays an important role in allowing axons to form synaptic connections. The pathway plays a role in growth cones and axons, rearranging the local cytoskeleton and plasma membrane[53]. Studies have found in ASD strong evidence implicating axon-guidance proteins in this disorder. ASD is largely caused by defects in neural development with the mechanism for this being explained by changes in axon guidance genes that impact neuronal morphology and structural connectivity during development. Changed expression or function of these genes contribute to the causes of ASD. Further explanation of this can be found when examining post-mortem brain tissue of patients with ASD which display decreased expression of axon guidance proteins. The general belief is ASD is caused by decreased connectivity of specific brain regions and increased connectivity of others which can be explained by axon guidance[76]. The significantly enriched GO terms for biological processes are axon guidance, negative regulation of axon extension, axon extension involved in axon guidance, and negative regulation of nervous system development. The listed biological GO terms are related to ASD as ASD is a disorder with neurodevelopmental causes and these GO terms relate to processes involved in neurodevelopment and axon guidance[76].

[IDP-25288-2.0] Ras signaling pathway was one of the leading pathways in KEGG analysis. The RAS-MAPK pathway serves many functions including mediating cellular responses to growth signals and activating several pathways which are involved in fundamental cell processes such as proliferation, differentiation, and adhesion[77]. Studies have shown that an attributing factor to the pathogenesis of ASD is dysregulation of the Ras MAPK signaling pathway. Looking specifically at studies of RASopathies that are a group of disorders caused by mutations of the Ras/MAPK pathway gene, an increased incidence of ASD has been noted in RASopathies. A study on the prevalence of Neurofibromatosis Type 1 a distinct RASopathy found high prevalence of ASD in NF1[78]. A study comparing affected probands who had various RASopathies (neurofibromatosis type 1 (NF1), Noonan syndrome (NS), Costello syndrome (CS), and cardio-facio-cutaneous syndrome (CFC)) and unaffected sibling controls found evidence of increased ASD traits in probands compared to controls and suggesting dysregulation of Ras/MAPK signaling during development may be implicated in ASD risk. The significantly enriched GO terms for biological processes are axonogenesis, regulation of cell morphogenesis, stem cell proliferation, cell growth, and developmental cell growth. The biological GO terms developmental cell growth is related to ASD as many of the risk genes associated with this disorder have to do with neurodevelopment and are involved early in brain development[79].

**Supplementary Notes III: Top SNVs' role in eQTLs regulating genes for neuropsychiatric disorders.**

***Schizophrenia*.**

For IDP-25190-2.0, three regulated genes, CD151 [AL049840.1], COL21A1[80, 81], and AL049840.1[82], have direct associations with SCZ, and PNPLA2 has been associated with other psychiatric disorders, including ADHD[83] and depression[84](**Figure 4A**). For IDP-25306-2.0, three out of five regulated genes, including XRCC4[85-87], TMEM167A[88], and WDR74[89], play an important role in the pathophysiology of SCZ . CENPK has also been related to mental disorders and cognitive impairments[90]. For IDP-25094-2.0, four regulated genes, NCKIPSD, CCDC71[91], GPX1[92, 93], and AMT[94-96], have direct associations with SCZ and other psychiatric disorders[91, 97, 98].

***Major Depression Disorder*.**

For IDP-25152-2.0, both AGMAT[99, 100] and PADI4[101, 102] have direct associations with MDD, while AJAP1[103] and PLEKHM2[83, 104] play an important role in the neuropsychiatric diseases, including BIP and SCZ (**Figure 4B**). For IDP-25905-2.0, MACROD2 is most commonly associated with ASD[105], although, it is also found to be related to ADHD[106], SCZ[107], and MDD[108]. Previous proteomics study has revealed that ACTR1A is involved in the anti-depressive mechanisms[109, 110]. MAP4K4, as an emerging therapeutic target in cancers[111, 112], has also been shown to be differentially expressed in BPD postmortem brains. For IDP-25862-2.0, ZNF84 has been found to be significantly dysregulated in the MDD patients[113, 114]. RP11-73M18[115] and APOPT1[116, 117] were also consistently implicated in both MDD and SCZ. For IDP-25270-2.0, FBXO42 is associated with the risk for MDD[118]. RP11-108M9.6 and RSG1 are both regulated genes underlying IDP-25284-2.0 for SCZ, which have been shown to be related to many psychiatric disorders[83, 84]. For IDP-25703-2.0, TTLL10 and TTLL10-AS1 are hypomethylated in Parkinson's disease[119] and annotated to the 58 CpG sites associated with educational attainment[120]. ATAD3A mutations can cause severe mitochondrial dysfunction[121] while mitochondrial dysfunction is a key player in the manifestation of depression[122]. Another study has shown that ATAD3A oligomerization promotes neuropathology and cognitive deficits in Alzheimer's disease models[123]. CNKSR1 interacts with proteins that have already been shown to be associated with intellectual disability (ID) and further proves to have an important role in brain development[124].

***Bipolar Disorder.***

The top five regulated genes that we identified for IDP-25906-2.0 are BLVRA, AFAP1L1, SLC44A5, RFTN1 and COA1 (**Figure 4C**). Among those five genes, SLC44A5[125] and RFTN1[126] had direct association with BPD. AFAP1L1[127] has been reported in patients with depression or depression-like symptoms. Given that BPD patients suffer from mood disorders, including feelings of depression and mania, AFAP1L1 is very likely to be associated with BPD. BLVRA, an enzyme that converts biliverdin to bilirubin, has recently emerged as a key regulator of the cellular redox cycle[128]. Bilirubin has been found negatively correlated with many neuropsychiatric and neurodegenerative disorders, including BPD, SCZ, Alzheimer's disease, dementia, etc[129-131]. BLVRA is involved in the pathogenesis of Alzheimer's disease[132-134].

***Autism spectrum disorder.***

For IDP-25869-2.0, TPRG1L is involved in the molecular pathways in ASD[135, 136]. Previous studies showed that the PIK3CD/p110δ catalytic subunit is one such isoform relevant for neurodevelopmental diseases, with PIK3CD being associated with increased risk for SCZ[137], and increased expression being identified in patients with SCZ or ASD[137-139]. Genetic variants within SLC25A12 gene have recently shown to be strongly associated with the etiology of ASD[140]. For IDP-25096-2.0, SLC16A8 is a known risk gene of glioma/glioblastomas[141]. In TWAS analysis, SLC16A8 was significantly associated with multiple white matter microstructure traits[142]. PICK1 mediates synaptic recruitment of AMPA Receptors at neurexin-induced postsynaptic sites, and mutations of human neurexin and neuroligin genes were found to be associated with ASD and mental retardation[143, 144]. CYFIP1[145] is also known as “specific Rac1-activated” (SRA1) protein[146]. The CYFIP1/SRA1 gene is located in a chromosomal region linked to various neurological disorders, including intellectual disability, ASD, and SCZ[147]. CYFIP1 is also close to a region critical for two ASD-related syndromes: the Angelman and Prader-Willi syndromes[148-150]. For IDP-25196-2.0, C1orf159 overlapped with CNV that are shown to be associated with ASD[151]. RP1-283E3.4 is up-regulated in POLR3B mutant patients[152], that implicated with intellectual disability[153]. PEX10 has been implicated in phenotypes of neurodevelopmental disorders, including Prader-Willi syndromes[154, 155], that are characterized by social difficulties that lie along the ASD[156]. CLCNKA are interacted with ASD-related genes in kidney and pituitary specific networks[157]. For IDP-25647-2.0, KCNAB2-depleted mice show significant deficits in associative learning and memory, suggesting a contributory role in the cognitive and neurological impairments observed in ASD patients[158-160]. H6PD, also referred to as glucose dehydrogenase (GDH), has extensive sequence similarity to G6PD and is believed to have been derived from this enzyme by sequence divergence following a duplication event[161, 162]. Metabolic alteration due to the deficiency of G6PD is associated with ASD. The G6PD deficiency increases ROS production which expedites neuroinflammation[163]. RSG1 is regulated genes underlying IDP-25284-2.0 for SCZ and IDP-25270-2.0 for MDD. The small GTPase RSG1 controls a final step in primary cilia initiation[164]. Previous studies showed that primary cilium acts as a cellular antenna sensing extracellular signals during brain development, and its formation is diminished in SCZ, ASD, BIP and MDD[165, 166]. For IDP-25288-2.0, DCAKD is recognized as playing a role in ASD[167]. NMT1 is essential for growth and development, during which rapid cellular proliferation is required, in a variety of organisms[168]. Deficiency in NMT1 exhibited overt growth, developmental, or behavioral defects[169]. SLC16A8 is associated with ADHD[170]. PICK1 mediates synaptic recruitment of AMPA Receptors at neurexin-induced postsynaptic sites, and mutations of human neurexin and neuroligin genes were found to be associated with ASD and mental retardation[143, 144](**Figure 4D**).

**Supplementary Notes IV: Running BrainXcan**

BrainXcan aims to analyze complex traits and their relationship with genetic markers of brain MRI-based characteristics, to identify important features that are specific to certain regions or applicable to the entire brain. BrainXcan takes GWAS summary statistics as input and return the association between GWAS phenotype and a list of brain image-derived phenotypes (IDPs).

The open-source implementation of BrainXcan is available at (https://github.com/hakyimlab/brainxcan ). We run BrainXcan using the default setting listed under the “Software documentation” section, and a detailed example can be found here: https://liangyy.github.io/brainxcan-docs/docs/example.html .

We first cloned BrainXcan code repository and installed all the BrainXcan software dependencies. To perform BrainXcan analysis, we need to prepare the following files:

1. The BrainXcan database and meta files, which was download from brainXcan database zenodo link: https://zenodo.org/record/4895174#.ZCO3snbMKMI
2. A GWAS summary statistics file, if there is no GWAS summary statistics file, we need to first generate summary statistics file from genotyping or sequencing data. The GWAS summary statistics file should contain information below:
   1) SNP rsID (we use rsID as SNP identifier) and chromosome of the SNP
   2) Effect and non-effect allele
   3) Group 1: effect size, standard error of effect size (zero-centered) or Group 2: z-score, GWAS sample size, allele frequency
3. Preparing the BrainXcan configuration file. Below is an example of the configuration file that we used for BrainXcan:

```
# ------------------------ Input/Output paths ------------------------------ #
# for the paths, please use full path to avoid issues
datadir: 'path-to-braixcan-database/brainxcan_data'
outdir: 'brainxcan_test'
prefix: brainxcam'
brainxcan_path: 'path-to-brainxcan-repository/brainxcan'
# ------------------------ GWAS formatting information --------------------- #

gwas: 'path-to-gwas-summary-statistics'
snpid: 'variant_id'
effect_allele: 'effect_allele'
non_effect_allele: 'non_effect_allele'
chr: 'chromosome'

# option 1 (high priority)
# Note: we assume effect is centered around zero, so use log(OR) for case control study
effect_size: 'effect_size'
effect_size_se: 'standard_error'

# option 2
zscore: 'zscore_2'
```

```
sample_size: 'sample_size'
allele_frequency: 'frequency'

# ------------------------- Generating BrainXcan region ---------------------- #
# if want to generate interactive html to present the regions
bxcan_region_vis: True # or False

# ------------------------- Optional: default values are listed -------------- #
# no need to specify if you'd like to go with default

# ancestry population of the gwas (used for MR). Options are populations in 1000G: AFR,
AMR, EAS, EUR, SAS
gwas_pop: 'EUR'

# IDP prediction model type: ridge or elastic_net
model_type: 'ridge'

# IDP sets to use: original or residual (after PC adjustment)
idp_type: 'residual'

# CV Spearman cutoff on models (only models passing this criteria will be shown)
spearman_cutoff: 0.1

# parameters to define signif BrainXcan results for MR
signif_pval: 1e-5
signif_max_idps: 10

# parameters used in defining instrument in MR
ld_clump_yaml: '{datadir}/mr/ld_clump.yaml'

# path to R and Python
rscript_exe: 'Rscript'
python_exe: 'python'
plink_exe: 'path-to-plink'

# generate empirical zscores with simulated weights in BrainXcan
bxcan_empirical_z: False
bxcan_empirical_z_seed: 1
bxcan_empirical_z_nrepeat: 1000

# generate empirical zscores with LD block-based permutation in BrainXcan
## path to the BED file (TAB-delimited base0 with header chr, start, stop)
bxcan_ldblock_perm: null
bxcan_ldblock_perm_seed: 1
bxcan_ldblock_perm_nrepeat: 10
```

4. Running BrainXcan pipeline. The general command is:

```
# conda activate brainxcan

REPO=path-to-brainxcan-repository
CONFIG=path-to-config-file

snakemake \
  -s $REPO/Snakefile \
  --configfile $CONFIG \
  SBrainXcanOnly
```

#### Supplementary Figures

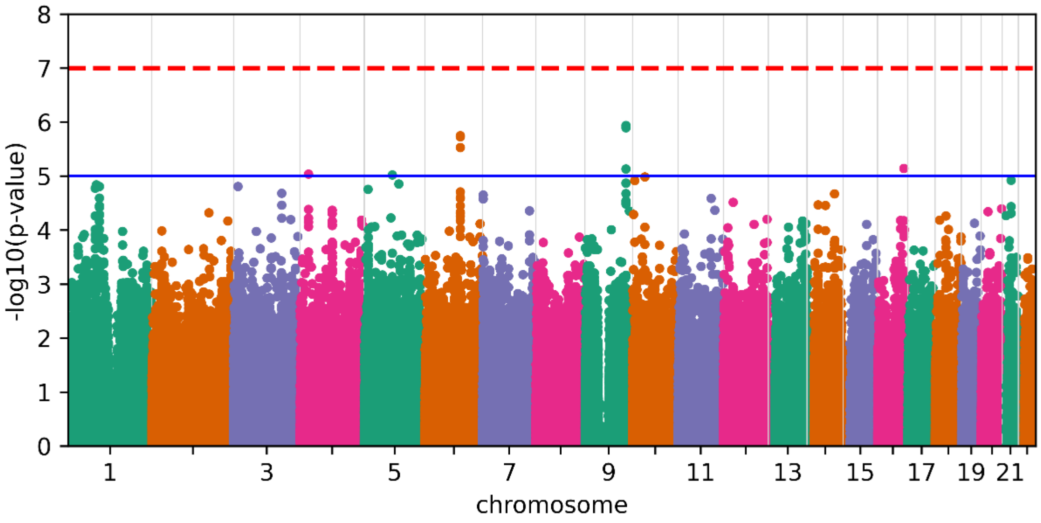

**Supplementary Figure 1. Manhattan plots of applying standard GWAS to Schizophrenia.** The stringent cutoff of Bonferroni correction is showed as the dash red line, and a loose cutoff of p-value=$10^{-5}$ is showed as a solid blue line.

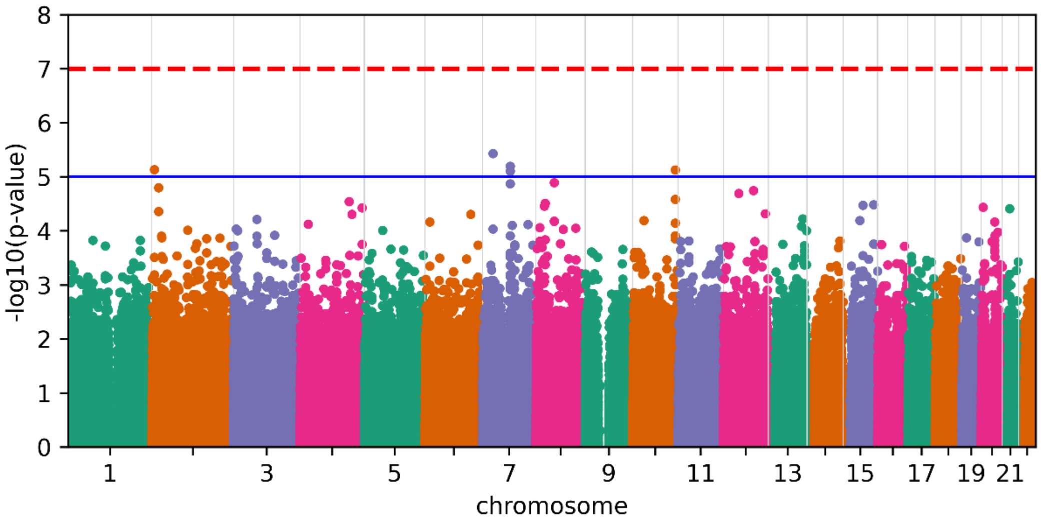

**Supplementary Figure 2. Manhattan plots of applying standard GWAS to Major Depression Disorder.** The stringent cutoff of Bonferroni correction is showed as the dash red line, and a loose cutoff of p-value=$10^{-5}$ is showed as a solid blue line.

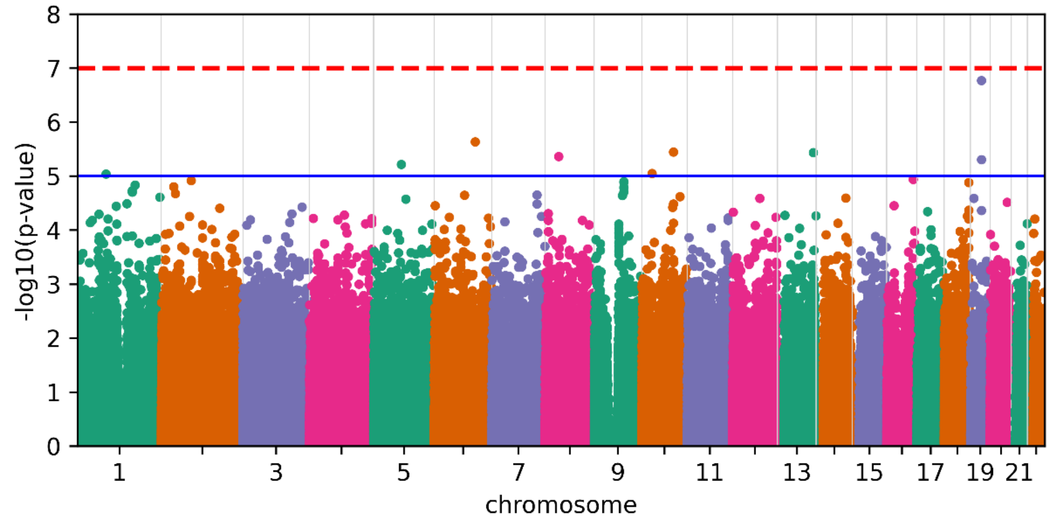

**Supplementary Figure 3. Manhattan plots of applying standard GWAS to Bipolar Disorder.** The stringent cutoff of Bonferroni correction is showed as the dash red line, and a loose cutoff of p-value=$10^{-5}$ is showed as a solid blue line.

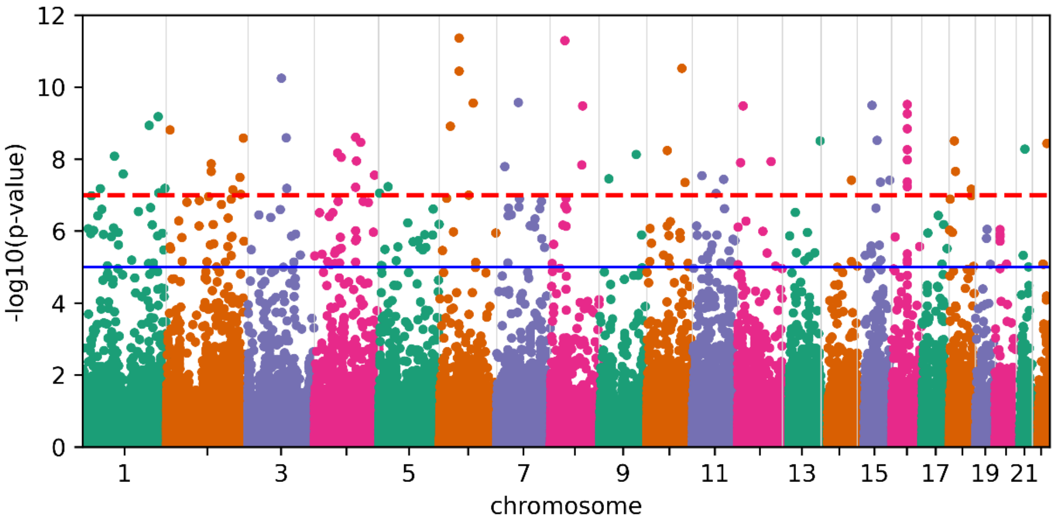

**Supplementary Figure 4. Manhattan plots of applying standard GWAS to Autism Spectrum Disorder.** The stringent cutoff of Bonferroni correction is showed as the dash red line, and a loose cutoff of p-value=$10^{-5}$ is showed as a solid blue line.

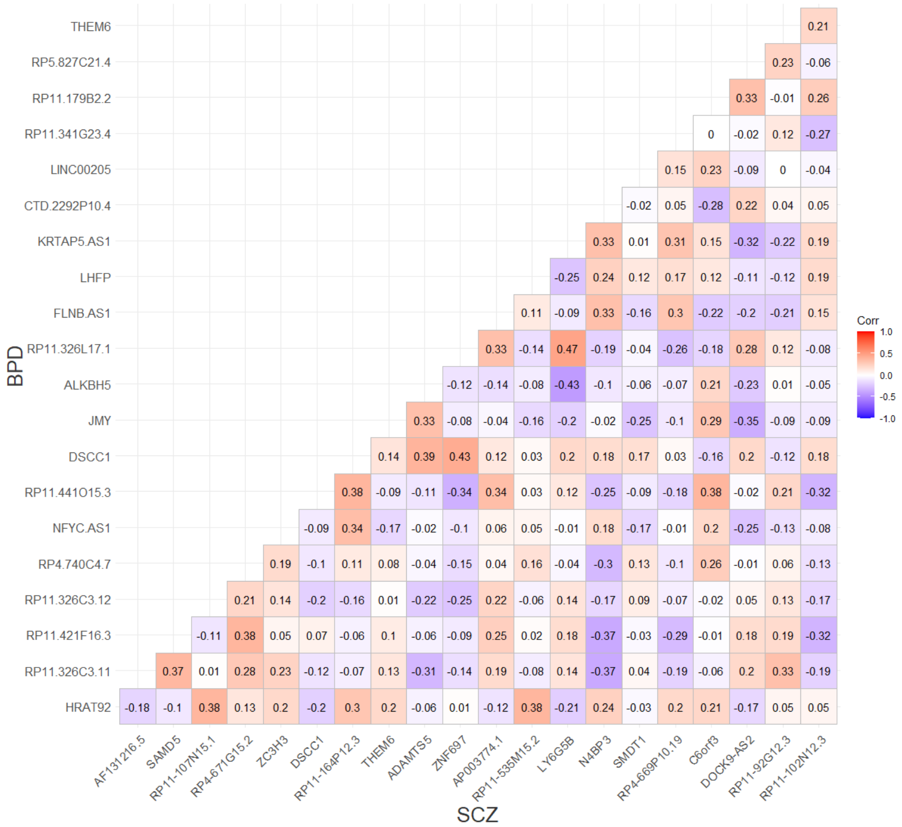

**Supplementary Figure 5.** A correlation matrix between the expression level of cerebellum eQTLs that were identified from BPD and SCZ.

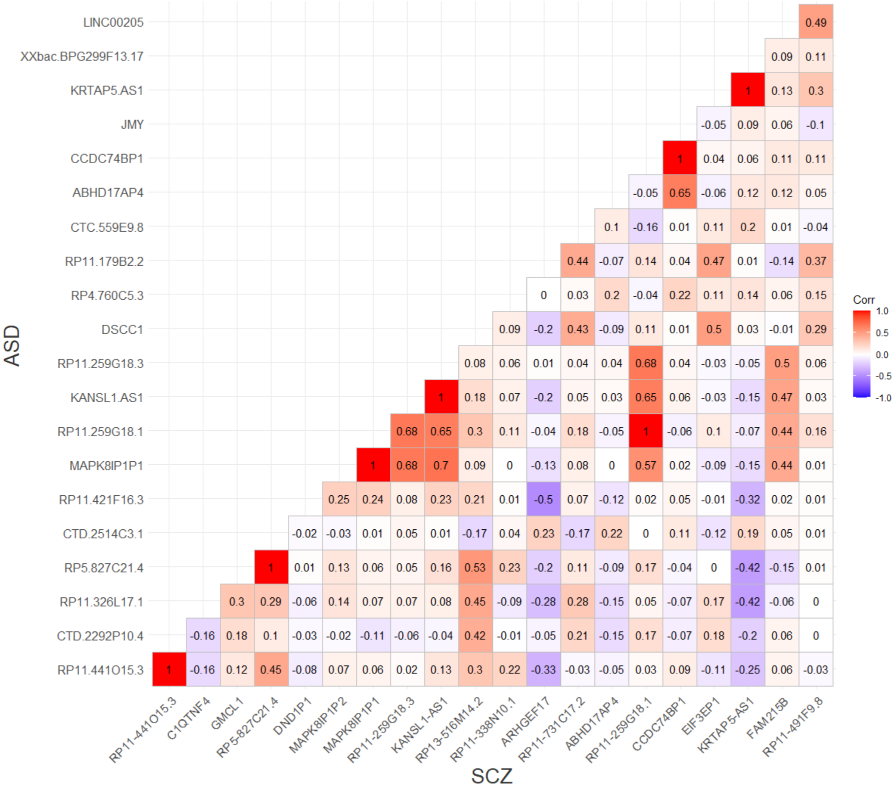

**Supplementary Figure 6.** A correlation matrix between the expression level of cerebellum eQTLs that were identified from ASD and SCZ.

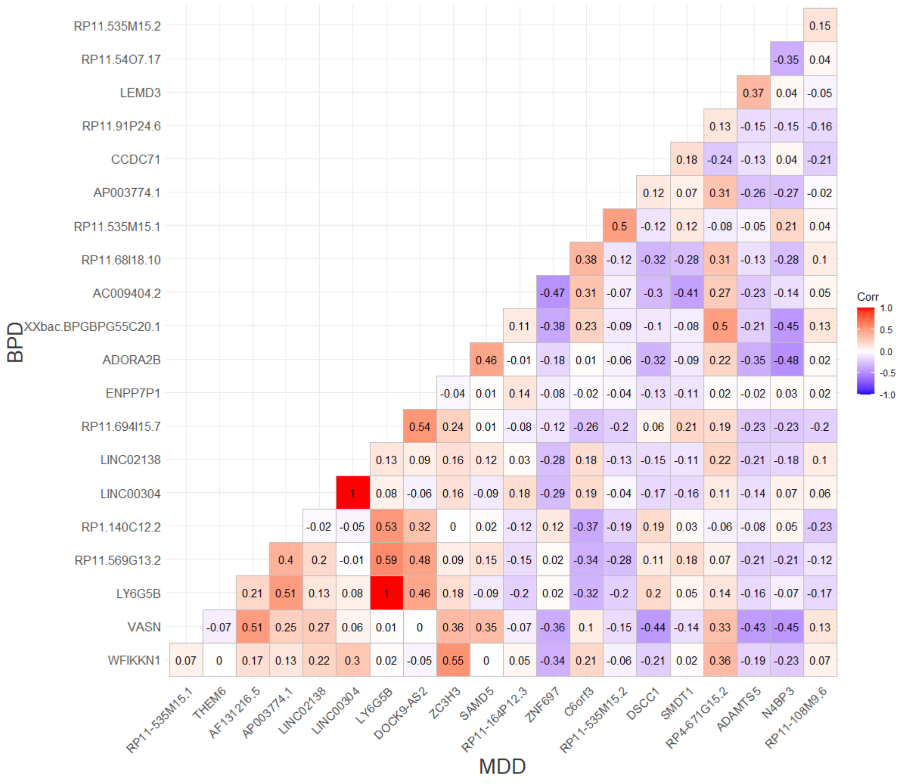

**Supplementary Figure 7.** A correlation matrix between the expression level of cerebellum eQTLs that were identified from BPD and MDD.

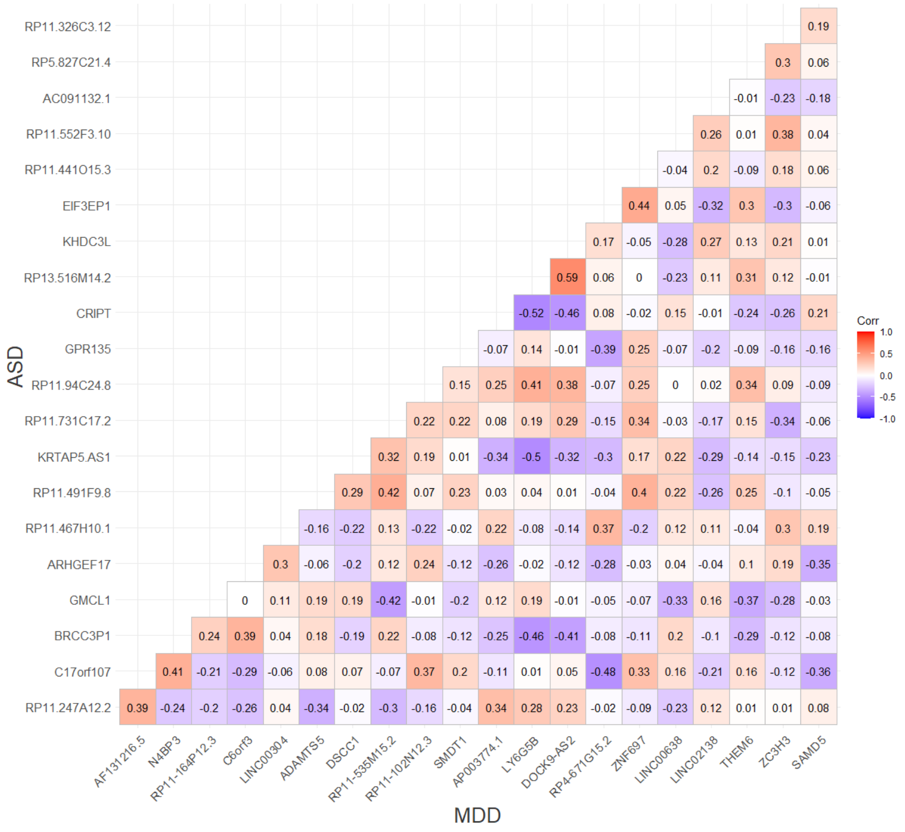

**Supplementary Figure 8.** A correlation matrix between the expression level of cerebellum eQTLs that were identified from ASD and MDD.

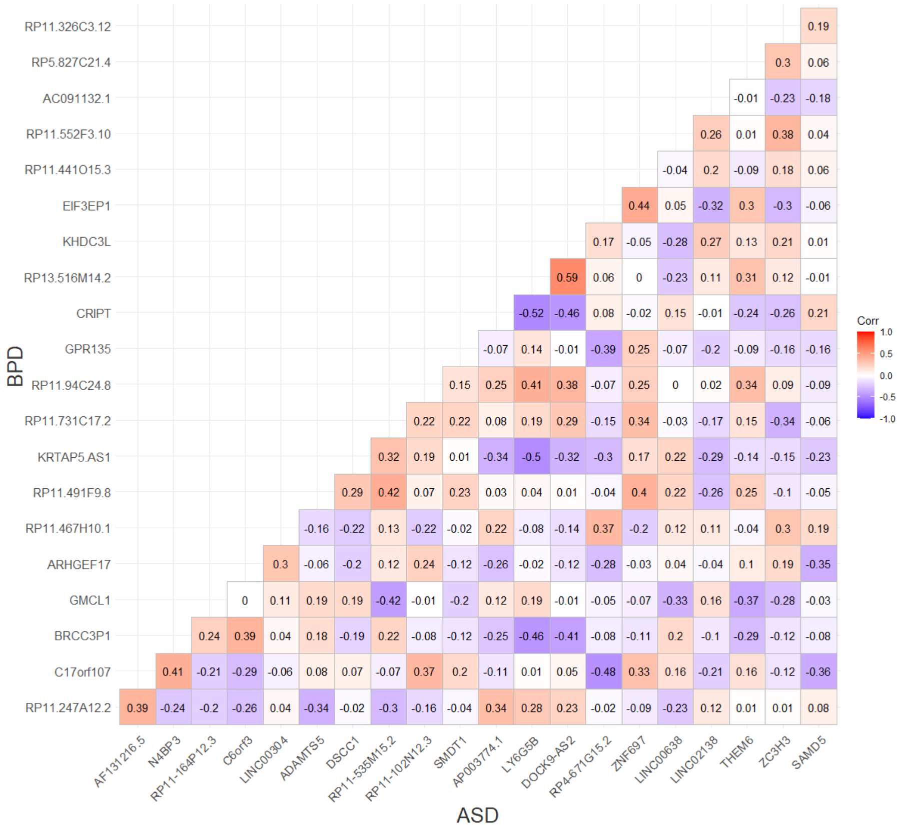

**Supplementary Figure 9.** A correlation matrix between the expression level of cerebellum eQTLs that were identified from ASD and BPD.

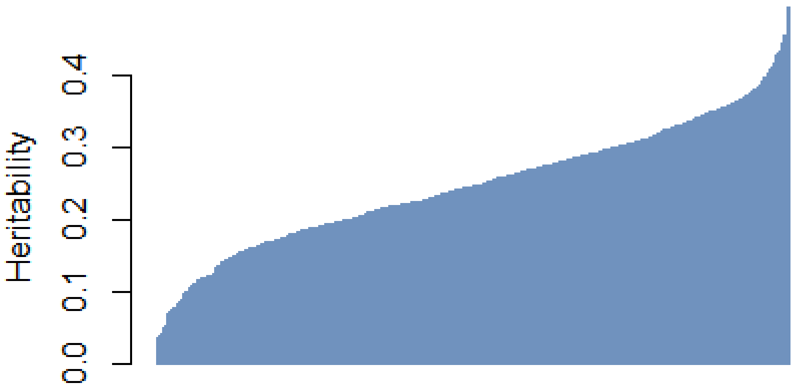

**Supplementary Figure 10. Heritability of each IDP using GCTA and the UK Biobank dataset.** The overall heritability is ranging from 0.02 to 0.49.

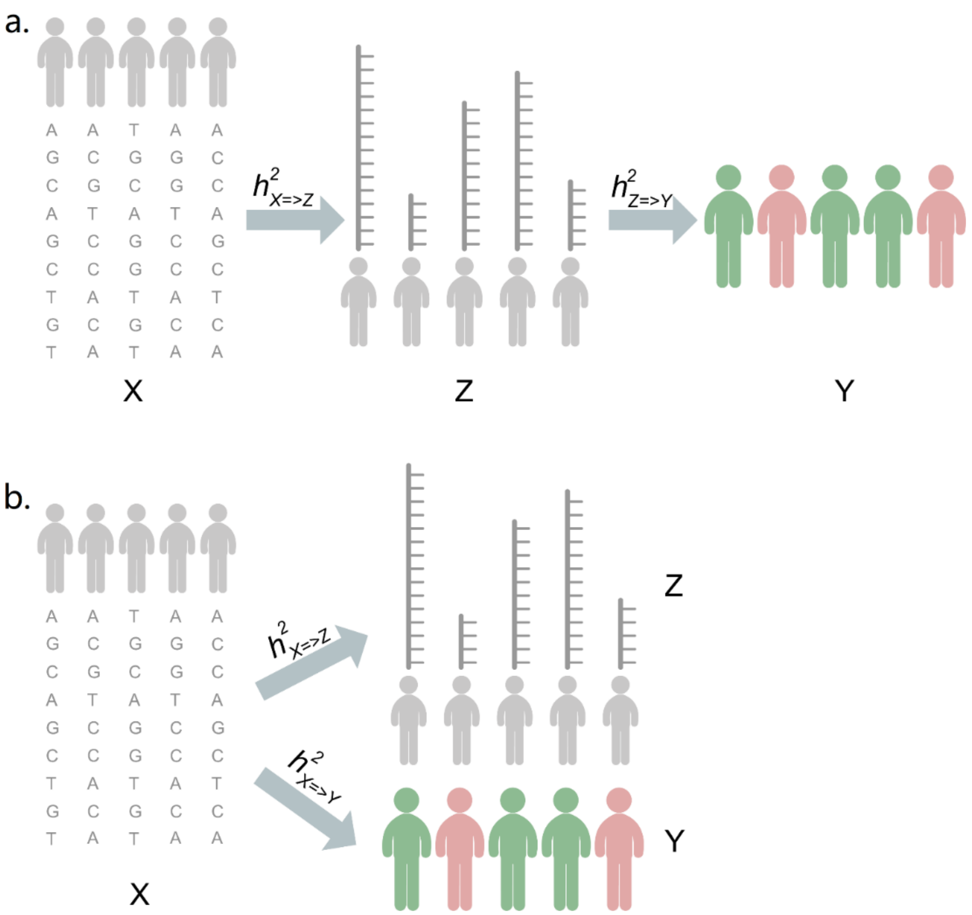

**Supplementary Figure 11.** Causality (a) and Pleiotropy (b) scenarios for genotype (X), image phenotype (Z) and phenotype (Y).
